# Personalised prediction of anomalous bone regeneration in multiple myeloma patients

**DOI:** 10.64898/2026.10.26.761863

**Authors:** Fiona G. Gibson, Rebecca E. Andrews, Michelle A. Lawson, Andrew D. Chantry, Margaret A. Paggiosi, Catherine Handforth, Janet E. Brown, Prash Thayaparan, Sean Molloy, Julia C. Shelton, Xinshan Li, Enrico Dall’Ara, Stefaan W. Verbruggen

## Abstract

Recent advances in immunotherapies have dramatically extended the lives of patients with multiple myeloma, resulting in a growing population of elderly patients suffering from damage to their vertebrae caused by this blood cancer. Clinicians typically intervene to fill in these lytic lesions in an attempt to reduce the risk of vertebral collapse and spinal cord injury, however observations of rapid bone regeneration post-cancer treatment suggest that surgery may not always be required. Here we develop a series of mechanistic computational models, optimised using cohorts of healthy volunteer and patient data, to show that vertebrae of recovering multiple myeloma patients rapidly strengthen in a manner not entirely explained by known bone remodelling mechanisms. Furthermore, using pathways informed by genomics data and patient-specific bone biomarkers, we demonstrate the potential for these models to provide personalised predictions of bone disease trajectory.

## Introduction

Multiple Myeloma (MM), a plasma cancer that arises in bone marrow and leads to bone destruction, represents around 2% of new cancers every year in the UK and the number of cases and deaths has more than doubled worldwide in the last 30 years^1^. Myeloma treatments have significantly improved over this period, increasing the 5-year survival rate for patients with MM to 50% and 44% in men and women, respectively^2,3^. Nevertheless, approximately 90% of patients with MM experience myeloma-induced bone disease (MIBD), and the majority are left with bone lesions post-treatment due to the destructive nature of this bone disease. Patients with MIBD are most commonly affected by spinal lesions (80-90%)^4^ with 34-36% of patients suffering from spinal fracture^5^. Both during and post-cancer treatment patients with unstable and weak spines require intervention to prevent vertebral collapse and spinal cord damage. Anti-resorptives (e.g. bisphosphonates and denosumab) and surgery are currently the most common interventions, although surgery is associated with increased morbidity due to the frailty of MM patients^6^.

In more recent years, orthopaedic bracing during the active cancer treatment has been presented as an alternative to surgery, providing structural stability. This takes advantage of the fact that MM-induced bone lesions can remineralise dramatically following treatment. Remineralisation after radiotherapy is detected in 48% of bone lesions^7^, and occurs in between 24% and 46% of patients^7,8^. Furthermore, Balducci et al. estimated that the time required to achieve remineralisation was around 6 months^8^. Moreover, bracing has been found to result in less pain alongside increase mineralisation and stability^9–12^, with evidence that the benefit of non-surgical treatment outweighed the risks associated with surgery^13^ if the bracing was implemented within six months of presenting^14^. However, while this phenomenon has been observed and reported through clinical computed tomography (CT) scans in individual case studies, and our recent work has shown increased evidence of repair^15^, we do not know whether this remineralisation results in greater bone strength or reduced fracture risk.

Bone is an exquisitely adaptive tissue, and in healthy conditions will adjust its mineral composition and structure in response to the mechanical loads it experiences^16,17^. A delicate balance of signalling exists between the three main bone cells involved in remodelling; osteoblasts that form bone, osteoclasts that resorb bone, and osteocytes that sense mechanical stimuli and in response direct the other cells to remodel bone tissue. In MIBD, these pathways are disrupted leading to an imbalance in bone remodelling^18^, bone destruction and an absence of bone formation, resulting in the development of lytic lesions within the bone^19^. After cancer treatment, most of the MM cells have been destroyed, which should allow the bone to return to normal remodelling^20^. Now left with large areas of destruction, the bone is much weaker and therefore under much higher strain when loaded. These conditions, in theory, should normally lead to bone formation as increased local deformation leads to greater mechanical stimulus sensed by osteocytes^21–23^. In fact, surprising clinical and experimental observations have shown bone formation can occur at faster rates than expected in recovering MM patients^9,11^, which gives rise to the question of whether this is driven solely by well-understood mechanobiological adaptation or if an additional biological mechanism is involved.

While clinical CT scans can characterise the effects of bone remodelling on bone mineral density (BMD) and its distribution, bone turnover biomarkers measured in patient blood capture a snapshot of the mechanobiological pathways in action. Biomarkers are widely used clinically to assess bone metabolism and estimate changes in fracture risk^24^, in conditions such as osteoporosis to monitor treatment efficacy and disease progression^25^. In MM, where bone remodelling is profoundly dysregulated, monitoring such biomarkers might help to elucidate the mechanisms driving mineralisation changes and to improve predictions of bone adaptation in MM vertebrae, ultimately informing more effective treatment strategies.

The aim of this study was to develop the first patient-specific predictions of vertebral bone regeneration in MM patients with established bone disease. Therefore, our objectives were to (1) determine whether bone strength is decreased in MM patients, and whether strength recovers in the vertebrae following a bracing regime; (2) develop and calibrate an adaptive computational model to determine whether remineralisation post-chemotherapy is governed by normal bone mechanobiology; and, (3) determine whether CT scans and blood bone biomarkers can be used to predict MM patient bone recovery and thereby inform surgical options.

## Materials and Methods

Our CT scans of MM patients demonstrate rapid mineralisation in the clinic, which can result in quite staggering changes in bone shapes, detected as increased intensity on CT scans and clearly visible in a reconstructed 3D computational model in Figure 1. These rapid changes in bone structure likely alter bone strength and may occur due to changes in mechanical load or changes in cell signalling due to cancer cells within the marrow microenvironment, though the contribution of each set of signals remains unknown.

**Fig. 1:**
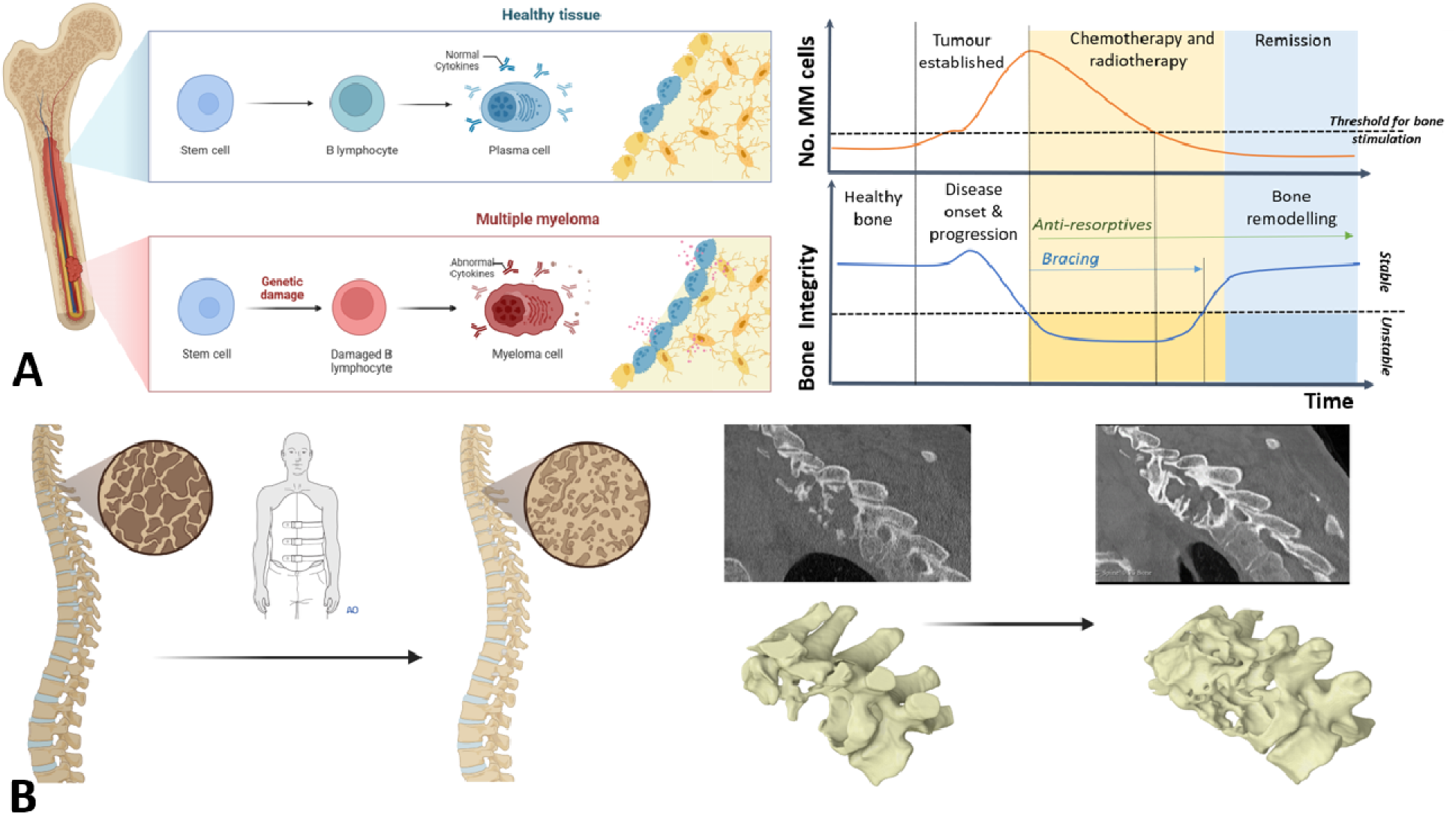
After treatment of multiple myeloma cells, bone rapidly remineralises with the application of bracing. (A) Schematic illustrating the development of multiple myeloma from plasma cells, and the typical disease and treatment trajectories. (B) Schematic demonstrating rapid remineralisation post-bracing, and a representative CT scan and resulting 3D reconstruction of rapid remineralisation of an osteolytic lesion in a multiple myeloma patient.

### Ethics approval

This study complies with all relevant ethical regulations for research involving human participants, with approval obtained for patient data from three distinct cohorts, as well as the computational analyses performed. Patient Cohort A: a retrospective collection of anonymised routine CT scans of MM patients from the Royal National Orthopaedic Hospital was approved by the Research and Innovation Centre at the RNOH (Reg. No. SE22.03). Patient Cohort B: Ethical approval for obtaining healthy volunteer CTs was part of the ANTELOPE clinical trial and was obtained from the South Yorkshire Research Ethics Committee in October 2016 (IRAS ID 206171)^26^. Patient Cohort C: Serial CT imaging and biomarker data were collected as part of the ‘Bone recovery after treatment study’^27^, which was reviewed by the Yorkshire & The Humber - Bradford Leeds Research Ethics Committee (REC 18/YH/0275) and approved by the Health Research Authority and Health and Care Research Wales (HCRW). Local ethical approval for the computational modelling performed on the anonymised patient scans from the above cohorts was obtained from both the University Research Ethics Committee at the University of Sheffield (Reference Number: 044189) and the Queen Mary Ethics of Research Committee at Queen Mary University of London (Reference Number: QME24.0767).

### Patient Data

CT scans of MM patients are not typically taken as part of standard clinical protocol, with MRI being the preferred imaging modality according to UK NICE guidelines^28^. Furthermore, quantitative CT (QCT) scans of healthy volunteers of the high resolution necessary to calibrate predictive computational models of bone remodelling are ethically challenging to gather. Therefore, for this study we sourced unique datasets of 3 patient cohorts (Figure 2): A) longitudinal clinical CT scans of braced MM patients; B) longitudinal densitometrically calibrated QCT scans of healthy volunteer age-matched inline phantom-calibrated controls; and C) longitudinal clinical CT scans of MM patients with matching longitudinal blood biomarker data.

**Fig. 2:**
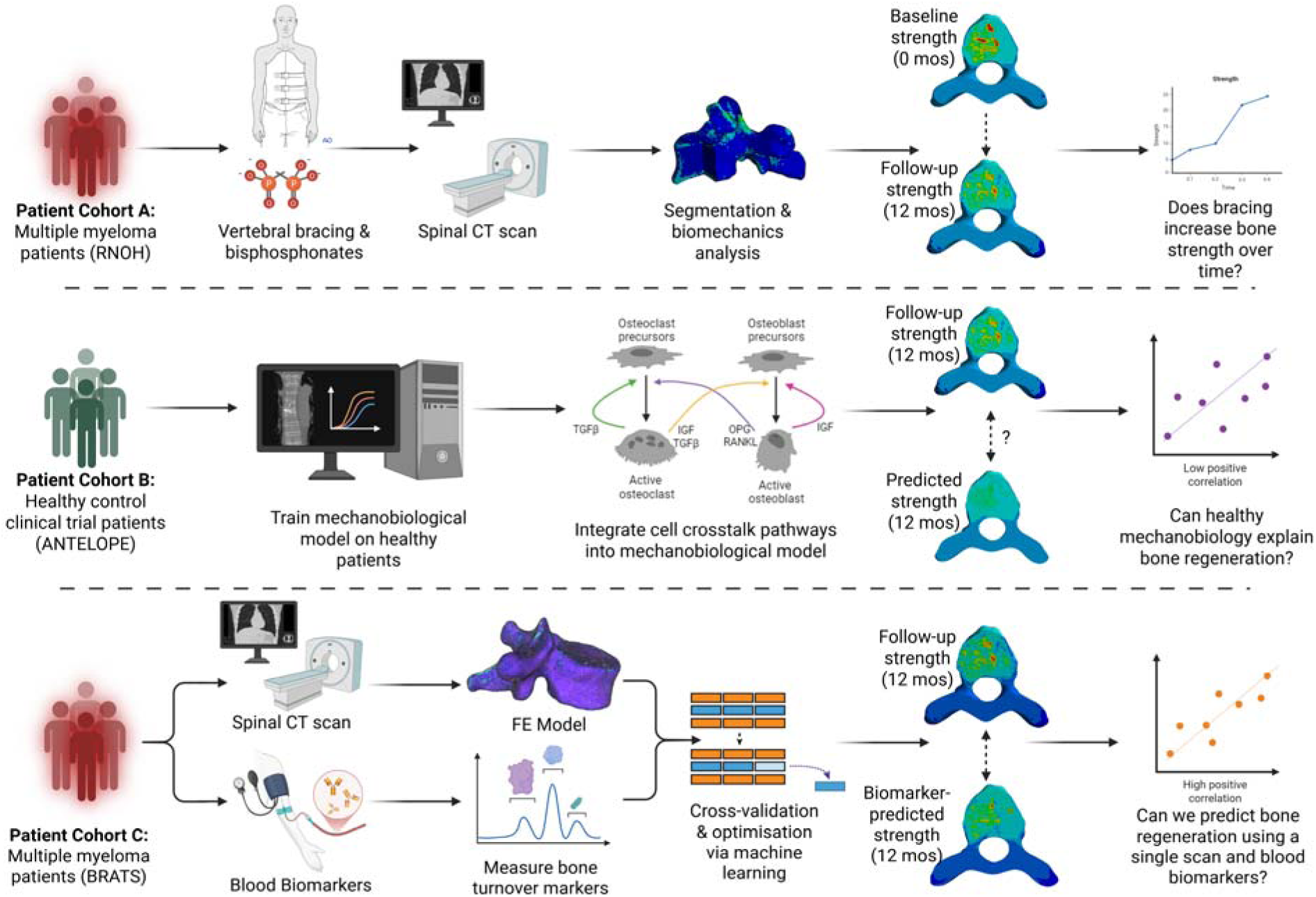
Three unique patient cohorts were used to investigate three research questions. Patient Cohort A comprised a timeseries of MM patients, investigating whether vertebral strength increased with remineralisation. Patient Cohort B was composed of non-cancer patients from a previous clinical trial, and was used to develop and optimise a mechanobiological algorithm of healthy bone remodelling. Patient Cohort C provided both CT scan and systemic blood biomarker data, allowing incorporation of patient-specific biochemical signalling pathways into model predictions.

### Computational Pipeline

#### Finite element modelling set-up

The finite element (FE) pipeline developed for all studies involved taking CT images, converting them into 3D models through segmentation, applying material properties and constraining the vertebral body for simulation to assess the mechanical properties (schematic of pipeline in Figure 3A). The FE pipeline has been used extensively to evaluate the mechanical properties of vertebrae with^29^ and without bone lesions^30–32^, and we have recently demonstrated its high intra- and inter-operator reproducibility^33^, assessed its predictive ability in predicting local bone formation^34^, employed it to identify critical bone metastatic lesions for vertebral strength^35^, and applied it to investigate the effects of adjuvant treatment on spinal fracture risk in prostate cancer patients^36^.

**Fig. 3:**
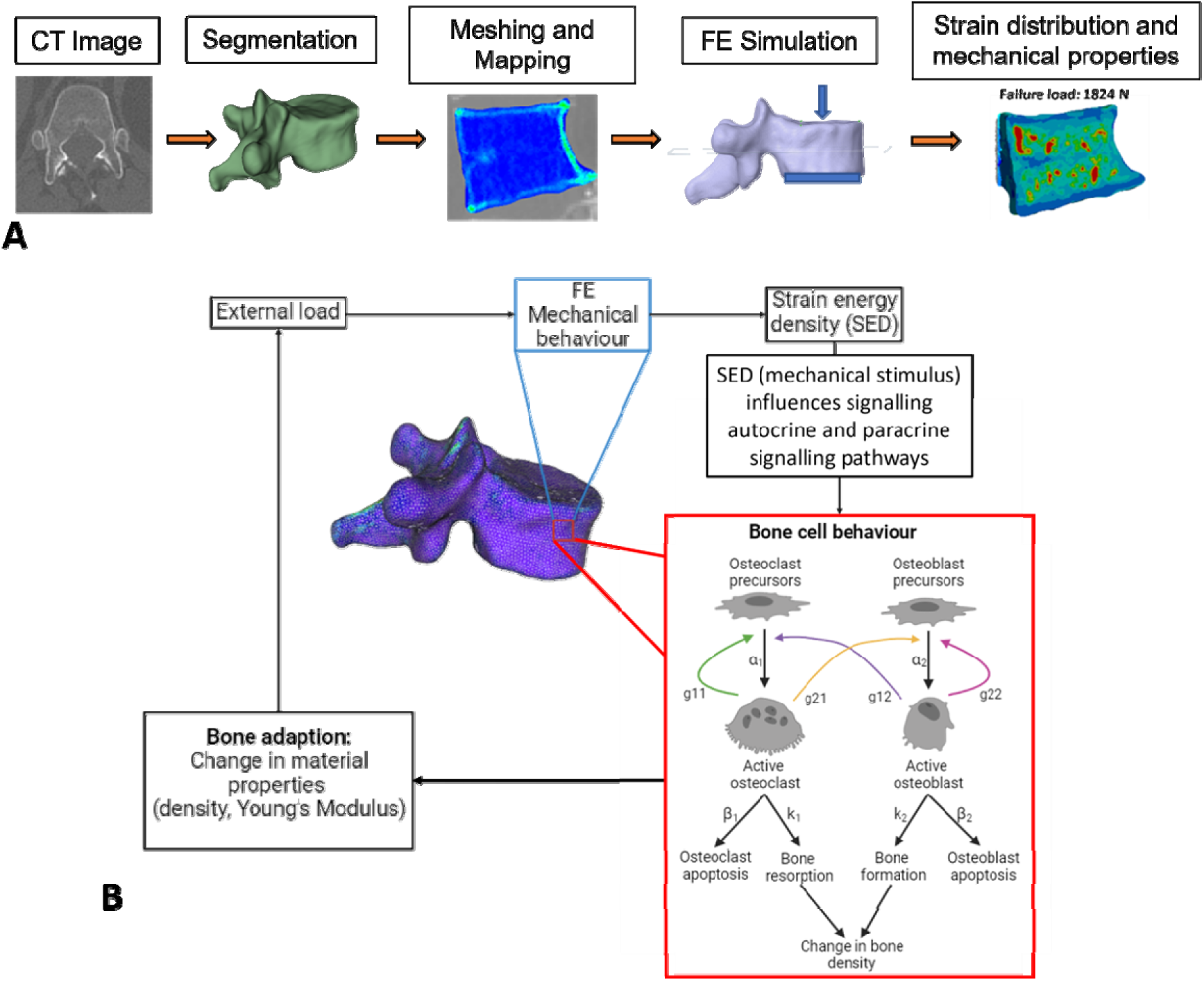
Computational pipeline used to build finite element model, develop mechanobiological model, and to train it to predict healthy bone remodelling. (A) Schematic showing the development of vertebral loading models, going from CT scan through segmentation and meshing to a finite element (FE) model. (B) Schematic explaining the steps taken in each iteration of the mechanobiological algorithm, illustrating how mechanical load is adapted to and how multiple parameters capture the cell-cell signalling contributions to the adaptation.

#### Verification, validation and Uncertainty Quantification

Given the extensive use of computational predictions in this study, robust verification, validation and uncertainty quantification (VVUQ) processes were employed to ensure robust and repeatable predictions. Verification was carried out, including mesh refinement using a protocol based on a previous verification study of metastatic vertebrae in other cancers^37^. These modelling protocols have been validated experimentally for vertebrae using digital volume correlation (DVC) by the authors^34^. Lastly, uncertainty quantification has been performed, as previously conducted by the authors^38–40^, using sensitivity analyses according to ASME guidelines for a research context of use^41^ and protocols from the EU CompBioMed project, of which Insigneo was a partner institute^42^.

Three additional specific verification procedures were carried out, as described in detail in the extended methods in the Supplementary Information: (1) an inter- and intra-operator variability study to ensure reproducibility of manual segmentation, (2) a comparison study to validate CT phantomless densitometric calibration against the gold standard of in-line phantom calibration^32^ and (3) a series of mesh convergence studies to verify mesh quality.

### Mechanobiological algorithm

#### Formulation

In order to investigate the underlying mechanisms for the rapid regeneration of bone observed in recovering MM patients, we must parse out what part of the remineralisation is driven by response to mechanical stimuli and the degree to which additional biological mechanisms are at play. In bone, mechanobiology is particularly important due to the hierarchical structure whereby mechanical loading at organ level influences the local mechanical stimuli and resulting biological response of bone cells and bone adaptation^43^. In MIBD, the cells and pathways involved are disrupted, leading to an imbalance in bone remodelling. While a long history of mechanobiological algorithms exists, dating back to Frost’s Mechanostat Theory^16^, there is currently no mechanobiological algorithm describing vertebral bone remodelling, in healthy volunteers or MM patients. Therefore, we developed an adaptive mechanobiological algorithm to investigate how mechanical loads affect biological processes.

In order to achieve this, the organ-level FE model described above was coupled with an algorithm representing the cell-level behaviours and interactions of bone cells using differential equations, which adjusted material properties based on biological pathways described by ordinary differential equations. The mechanical stimulus was defined based on the difference between a strain energy density (SED) in the FE model and a baseline reference stimulus^44^. The magnitude and nature of the mechanical stimulus (bone anabolic or catabolic) influenced the interaction between bone cells and the ratio between formation and resorption, essentially acting as a dimmer switch on the mechanoregulation (see Supplementary Information for more detailed formulation).

Briefly, the mechanical stimulus (*S*) was defined based on the difference between the ratio of SED to density (SED/ρ) calculated in the FE model and the reference stimulus from literature^21^. If the SED/ρ was above this reference value, the mechanical stimulus the element received was positive and if the SED/ρ was below this reference value, the mechanical stimulus the element received was negative.

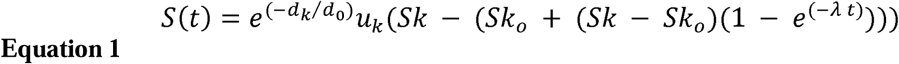

where *d_k_* = *OCd/N_rem_* Osteocyte density/number of remodelling sites, *d_0_* = osteocyte network connectivity, *S*(*t*) = mechanical stimulus, *u_k_* = osteocyte mechanosensitivity, *Sk* = SED/ρ per element, *Sk*_0_ = reference SED/ρ, *λ* = velocity of adaptation.

This mechanical stimulus influenced the number and activity of bone cells. The number of active osteoblasts and osteoclasts was then calculated by taking the steady state populations away from the updated population for that timestep (affected by the mechanical stimulus)^45^.

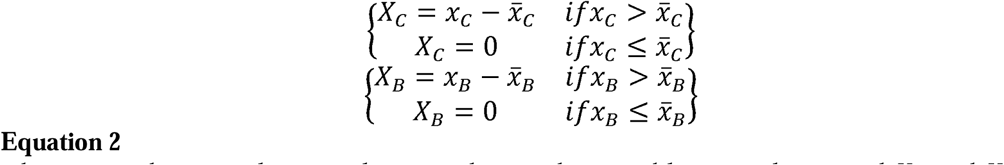

where *x_c_* is the osteoclast population and *x_B_* is the osteoblast population and *X_C_* and *X_B_* are the active populations of osteoclasts and osteoblasts, respectively.

The density of each element in the model was then updated based on the number of active osteoblasts and osteoclasts and the rate of bone resorption (k1) and bone formation (k2) respectively^45^.

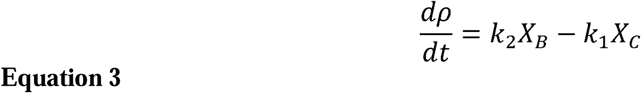

This mechanical stimulus input influenced the number of active osteoblasts and osteoclasts, but their activity could also be influenced by specific biological signalling pathways. Additionally, osteocyte density, osteocyte network connectivity and mechanosensitivity were also included in the formulation, due to their known important role in regulating mechano-adaptation, and were kept homogenous across the model in all studies.

### Modelled Biochemical Mechanisms and Gene Expression Database Analyses

A number of key pathways have been included in this algorithm that are known to play important roles in bone cell interactions and the bone remodelling cycle. A high mechanical stimulus reduced the expression of RANKL and increases OPG, reducing osteoclast differentiation and activity^46^. There were also signals from the bone matrix when bone is under mechanical stress - TGF-β signalling, which is sequestered within the bone matrix and released during bone degradation, promotes osteoblast differentiation and impedes osteoclast differentiation^47^. The combination of these pathways ensures normal bone remodelling and bone homeostasis. We grouped these signals, along with IGF, into two autocrine and paracrine pathways due to their known importance in bone remodelling in the context of MM^31^.

In MM, these pathways are disrupted leading to an imbalance in bone remodelling. The RANKL/OPG signal is upregulated whilst the Wnt signal is downregulated, increasing osteoclast activity and decreasing osteoblast activity respectively^18^. Furthermore, previous studies by our groups have shown that TGF-β inhibition could enhance bone repair in a mouse model of MM^48,49^. The combination of bone destruction and no new bone formation leads to the development of lytic lesions within the bone^19^. Additionally, our analysis of datasets in The Cancer Genome Atlas showed that these specific pathways are highly associated with significantly different survival outcomes in MM patients. Kaplan-Meir plots of recurrence-free survival in patients with high or low expression of genes encoding for key bone remodelling pathways (OPG, RANKL, TGF-β, IGF) were generated from The Cancer Genome Atlas (TCGA) using KMPlotter^50^. Analysis was performed on the Pan-Cancer DNA repositories, using n=980 MM samples.

### Sensitivity analysis of predictive algorithm

To quantify the uncertainty and sensitivity of the parameters within the mechanobiological algorithm, a sensitivity analysis was conducted on all parameters to determine which were most critical. Each of the 20 parameters was altered by ±50% and the local change in density was evaluated then averaged across all elements. This resulted in six specific parameters being selected as sensitive enough to require optimisation on a patient-by-patient basis. Additionally, all six selected parameters (k1, k2, α1, α2, β1 and β2) are directly related to key bone biomarkers that can be measured clinically (defined in Supplementary Information).

### Optimisation of mechanobiological algorithm parameters

To optimise the mechanobiological parameters, a MatLab script was developed which began by defining an objective function that ran the original mechanobiological algorithm, computing the simulated BMD and then calculating the standard error between the simulated density and the density measured in the follow-up scans. This function performed the optimisation to minimise the standard error by adjusting the parameters until convergence. Convergence was met when either the optimality tolerance, the standard error was below 1×10^-6^, or the step tolerance was below 1×10^-10^, where further changes in the input parameters did not significantly reduce standard error.

### Clinical bone biomarkers

A unique dataset of seven MM patients were collected as part of a pilot clinical study (BRATS), curtailed by the COVID-19 pandemic, which collected biomarker data alongside baseline and follow-up (12 months) CT scans^15^. The study protocol is explained on medRxiv^27^, and the patient data is detailed in Table S3, where the vertebrae were split into either ‘no lesion’, ‘small lesion’ where the lesion is <50% of the vertebral body and ‘large lesion’ where the lesion is >50% of the vertebral body. Biomarker data for procollagen type I N-terminal propeptide (P1NP), carboxy-terminal collagen-crosslinks (CTX) and sclerostin was obtained for 5 patients (Table S3) at 1, 2 and 3 months after baseline (for detailed description see Supplementary information).

P1NP is a known biomarker of bone formation, and is released by osteoblasts when they produce type 1 collagen to form the bone extracellular matrix^51^. In contrast, CTX is a specific and sensitive biomarker of bone resorption^25^ released when the collagen matrix is degraded by osteoclasts^52^. CTX and P1NP have both been recommended by the International Federation of Clinical Chemistry and Laboratory Medicine (IFCC) and the International Osteoporosis Foundation (IOF) in studies evaluating fracture risk assessment and treatment response^24^ and have been deployed in the context of MM^53^.

Sclerostin is known to be another indicator of bone turnover. Sclerostin is secreted by osteocytes and inhibits the differentiation and proliferation of osteoblasts^54^ by blocking the Wnt pathway^55^. While less commonly used in the clinic to monitor bone turnover than CTX and P1NP, sclerostin levels have been shown clinically to reflect inhibition of bone formation^56^.

The mechanobiological algorithm was applied to this dataset, adjusting the six mechanobiological parameters (k_1_, k_2_, α_1_, α_2_, β_1_, β_2_) for each patient. Then, a multiple linear regression analysis using leave one out cross validation (LOOCV)^57^ was performed in MATLAB (R2024a). The multiple linear regression assessed the relationships between the three biomarkers and each of the optimised parameters (k_1_, k_2_, α_1_, α_2_, β_1_, β_2_), as well as between the three biomarkers and both BMD and failure load (FL) at 1, 2, and 3 months after baseline, separately. LOOCV used all but one patient to train the algorithm and then tests on the final patient, increasing the training dataset for small cohort sizes^57^. It repeated this for all patients, increasing the robustness of the relationship compared to a simple linear regression. Following this analysis, parameters with a strong positive correlation between follow-up and predicted (r > 0.8) were then estimated using the linear regression equations utilising the biomarkers (Equation 1). All other parameters that had a negative, moderate or weak correlation (-1 < r < 0.8) were averaged across the cohort.

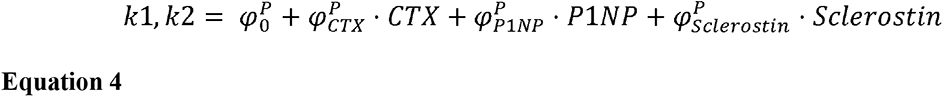

where, k_1_, k_2_ is the parameter to be predicted (also, α_1_, α_2_, β_1_, β_2_), 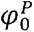 is the intercept, 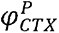 is the proportionality coefficient for CTX, 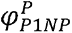 is the proportionality coefficient for P1NP, 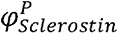 is the proportionality coefficient for Sclerostin.

## Statistical Analysis

Statistical analyses were preformed using Matlab, and are as described in figure legends.

## Results

### Multiple myeloma patients with large osteolytic lesions, increase rapidly in strength with bracing treatment

The computational models of MM patients with large lytic lesions (size >50% of the vertebral body) in Patient Cohort A significantly remineralised and strengthened over time, with lower strain observed in the vertebrae in follow-up CT scans (Figure 4). This was clearly shown as increased predicted failure load and ultimate strength. In contrast, for vertebrae without metastatic lesions or with small lesions (size <50% of the vertebral body) there was no obvious increase in BMD or mechanical properties evaluated at follow-up. This therefore confirmed our hypothesis that the previous clinical observations of rapid remineralisation following bracing treatment for MM patients did in fact lead to stronger vertebrae. However, the rapid changes observed in BMD and strength, particularly in vertebrae with large lesions, are not well understood; whether these changes are driven through normal remodelling or whether there was an additional biological mechanism influenced by the myeloma cells was still unknown.

**Fig. 4:**
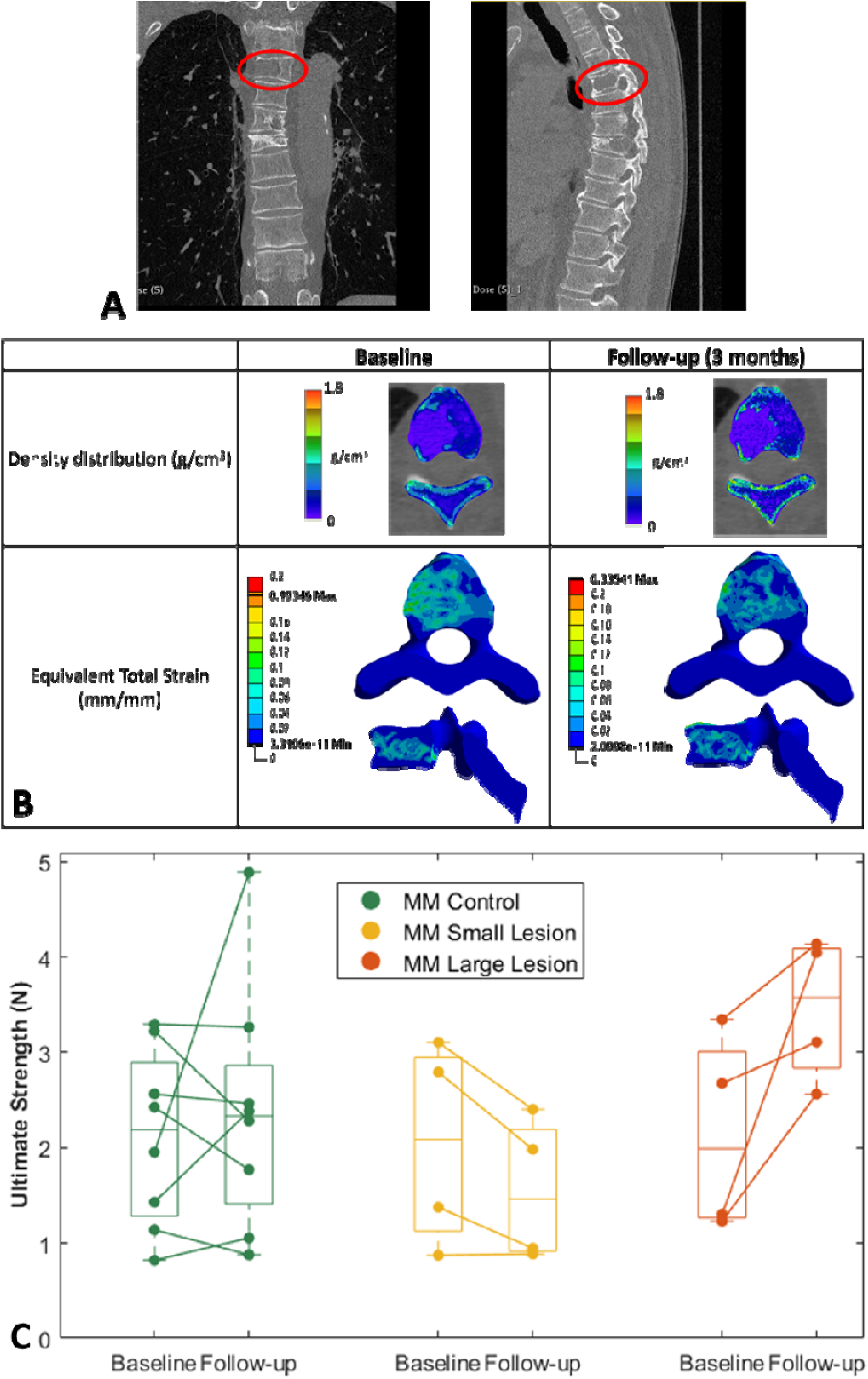
Large lytic vertebral lesions rapidly remineralise and strengthen with bracing post multiple myeloma treatment. (A) A baseline and follow-up scan of the same patient, with notable increases in BMD observed. (B) Representative scan with a large lesion showing marked increased in BMD and mechanical properties over time, resulting in decreased strain and fracture risk. (C) Large lesions demonstrate increased failure strength after bracing, but only in the follow-up scans of the vertebrae with large lesions.

### Mechanobiological algorithms cannot predict the rapid strengthening of vertebrae in recovering multiple myeloma patients, indicating dysregulation of signalling pathways

We next built a mechanobiological algorithm, a series of equations that govern how much mineral is generated or removed by cells in a specific region, in response to the mechanical stimulus they sense (Figure 3). We then optimised the parameters of the algorithm by training it on healthy age-matched volunteers so that it could predict healthy bone remodelling over 12 months when given a baseline CT scan.

Having coupled the organ-scale bone adaptation to cell-level behaviours of bone cells influenced by mechanical stimulus, the model was able to predict the changes in BMD and mechanical properties in the healthy volunteers (average percentage difference BMD: 2.13 ±2.36%, average percentage difference in failure load (FL): 3.64 ±3.72%; Figure 5). The model also predicted the BMD well for the MM vertebrae with small lesions (average percentage difference BMD: 3.49 ±4.63%; Figure 5).

**Fig. 5:**
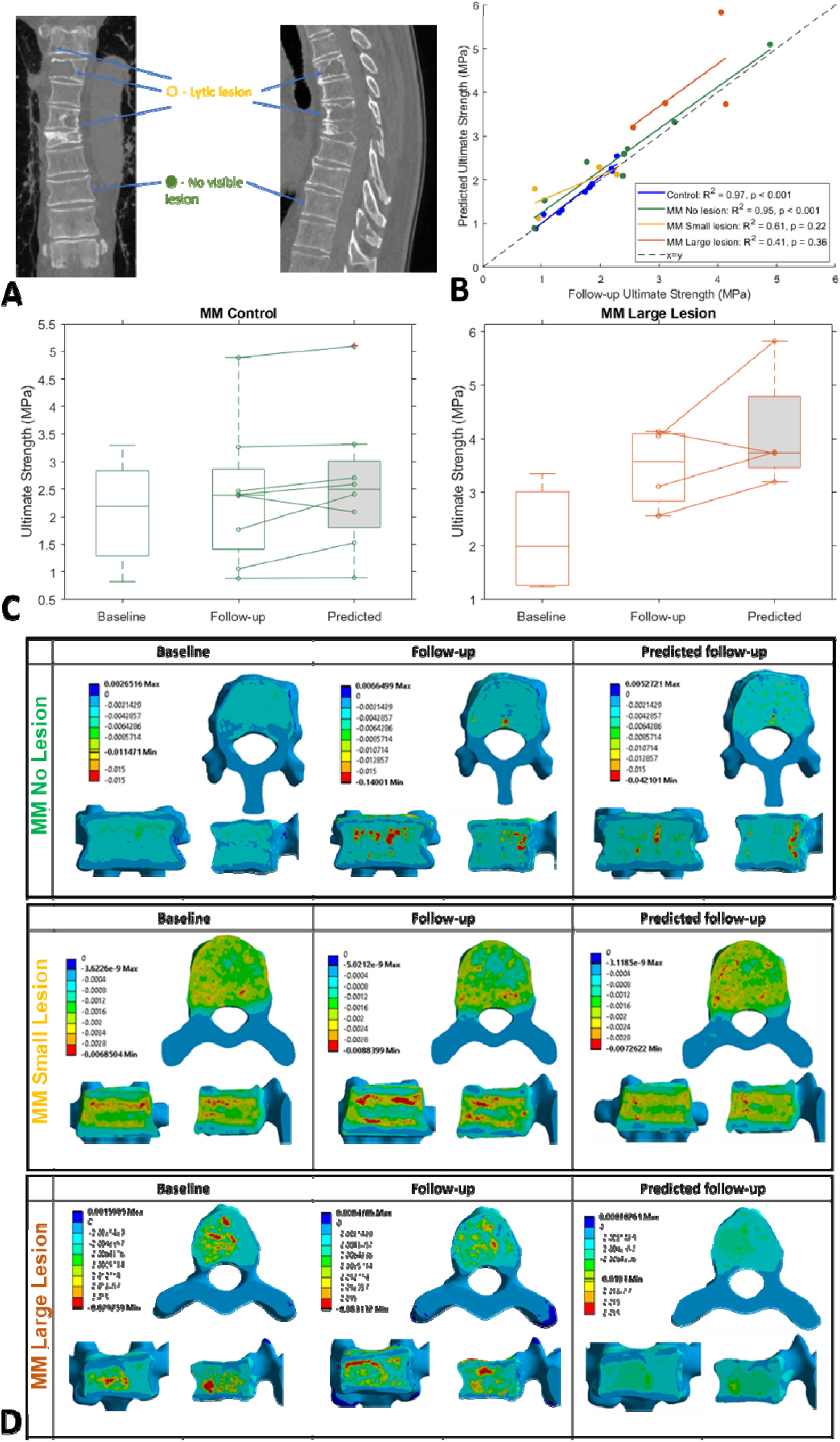
Mechanobiological algorithm trained on clinical trial data can predict changes in strength of healthy vertebrae and in vertebrae with small lytic lesions, but not in those with large lytic lesions. (A) Representative scan showing lesion-free vertebrae used as in-patient control vertebrae. (B) Mechanobiological algorithm applied to control vertebrae showed excellent predictions of follow-up vertebral failure strength despite being trained on a separate healthy dataset, but did not successfully predict strength in vertebrae with MM lesions. (C) Patient specific predictions of vertebral strength were highly consistent in control patients, but not in patients with large lesions; (D) Representative model predictions of compressive strain distributions in a control vertebra, a vertebra with small lesions, and a vertebra with large lesions, clearly demonstrating that the mechanobiological algorithm cannot predict changes observed in large lesions.

However, in vertebrae with large lesions, the predicted strain distribution does not accurately capture the observed changes in strain. This is particularly clear in the lesion region, where the compressive strains are much lower in the Predicted model compared to the actual Follow-up. Our algorithm was over-predicting bone regeneration in these vertebrae, strongly indicating that biological signals additional to mechanical stimuli were at play.

### Blood biomarker-informed mechanobiological algorithms augment patient-specific predictions of bone recovery

Lastly, having found that remineralisation in MM patients post-cancer treatments cannot be explained by healthy bone remodelling, we applied our novel algorithm to a new dataset in which we could adjust the biological parameters based on patient specific clinical biomarkers. By utilising bone biomarkers present in blood, we present the possibility of applying our mechanobiological bone adaptation algorithm to predict disease trajectory without the need for a follow-up CT scan. These updated models successfully predicted the BMD and failure load in three out of the five patients (average percentage difference BMD: 3.95 ±3.27%, FL: 7.1 ±8.5%; Figure 6), offering valuable insights into how early mechanobiological responses affect mechanical properties after 12 months. The other two patients are less well predicted, and this may be due to the presence of new lesions in the follow-up scan, indicating recurrence of MM. These models deploy the first biomarker-informed mechanobiological algorithm to predict patient-specific bone changes in MM, and are a promising step towards understanding and predicting patient trajectory and fracture risk in MM-induced bone disease.

**Fig. 6:**
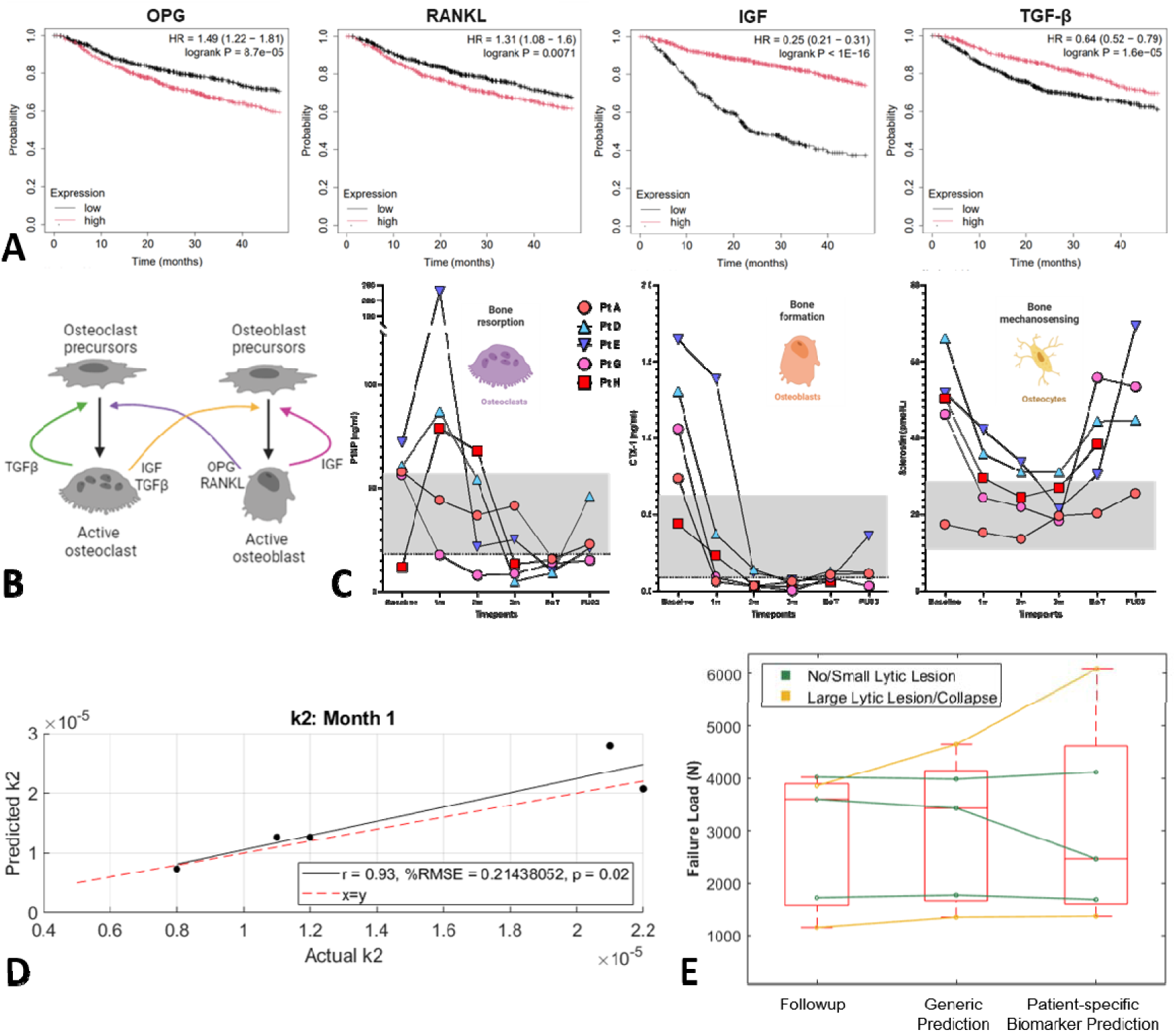
Incorporation of blood biomarkers improves patient-specific prediction of vertebral strength. (A) Kaplan-Meir plots of recurrence-free survival in patients with high or low expression of the genes encoding for bone related pathways in MM patients, generated from the Cancer Genome Atlas (TCGA) using KMPlot^50^. (B) Implementation of these signalling pathways into our mechanobiological algorithm. (C) Changes in measurements of key clinical blood biomarkers for bone turnover for resorption by osteoclasts (CTX), bone deposition by osteoblasts (P1NP) and mechanosensation by osteocytes (sclerostin)^15^. (D) Correlation from optimisation study showing that, when the algorithm is trained on biomarkers at 1 month after baseline a significant correlation exists. (E) Initial predictions demonstrate potential of biomarker-trained algorithm to predict bone regeneration 12 months after an initial CT scan.

## Discussion

This study shows for the first time that vertebrae severely damaged by MIBD rapidly regain their strength after first line cancer treatment, in a manner that may preclude the need for invasive surgery in some clinical cases. By developing computational models trained on a gold-standard clinical trial CT dataset of healthy vertebrae, we have also unveiled that this rapid remineralisation is not driven by normal mechano-adaptation mechanisms alone. Finally, by adapting our algorithm through the first reported dataset of CT scans and blood biomarkers from MM patients^15,27^, we have demonstrated the intriguing potential for novel improved predictions of bone remodelling to inform patient treatment and surgical options.

While robust VVUQ tests have been applied to all models developed in this study, the development of these models required a number of assumptions. One key limitation is the variability inherent in clinical CT scan quality, with clinical CTs acquired using different scanners, scanning protocols and resolutions, with this likely having some effect on the segmentation reproducibility. A more extensive study involving a large cohort of scans with varying protocols is necessary to assess the magnitude of this effect. This also applies to the effect of segmentation reproducibility on FE-predicted mechanical properties, as FE analyses depend on the accuracy of segmented geometries, meaning that inconsistencies or errors in segmentation could propagate through the pipeline and influence mechanical property predictions. Lastly, only one standard loading condition was tested within this FE framework, focusing on uniaxial compression as the standard mode of physiological loading. Additional work is necessary to investigate additional loading conditions, such as torsion, bending, and eccentric compression, in order to provide a more comprehensive understanding of how lytic vertebrae respond to different loading or fall conditions in everyday scenarios.

By employing a robust finite element pipeline, we have shown that significant changes in the mechanical properties of vertebrae with lytic lesions following treatment with orthopaedic bracing occur. The results demonstrated that over time, with remineralisation, the mechanical properties of the large lytic vertebrae improved, confirming clinical observations that remineralisation following bracing treatment for MM patients leads to an increase in vertebral strength. Indeed, there is early evidence within our BRATS study that rapid remineralisation occurs even without bracing^15^. In contrast, vertebrae with smaller lesions displayed either no change or a decrease in mechanical strength. This provides key evidence for clinicians that, in some cases, bone of patients will recover without the need for surgical intervention and the associated risk of infection.

Understanding how and why this remineralisation occurs in certain regions, whether it is purely mechanically driven or there are additional biological mechanisms involved, is key to applying this knowledge in the clinic. Using our newly developed mechanobiological algorithm, trained on clinical trial datasets and optimised using parameter optimisation techniques, we were able to use one CT scan to give a robust prediction of the bone shape and strength for both healthy volunteers and MM patient vertebrae with no lesions that was observed clinically at follow-up. Interestingly, the algorithm was unable to capture the rapid remineralisation evident in large lytic lesions, even after tuning the model to optimise parameters for specific patients. This confirmed our hypothesis that normal mechanobiology principles cannot explain the bone changes in MM patients’ vertebrae treated non-surgically, implying that additional biological mechanisms are at play in this recovery from MM. Indeed, this clinical question, along with is implications for pain, is being pursued as part of an ongoing clinical trial (BACBONE, NIHR175280)

By incorporating bone turnover markers that can be gathered clinically, we have unpicked the underlying mechanisms of how disruption of bone biochemistry by MM cells can lead to structural changes in bone. Applying this knowledge in our novel algorithm, we have also provided the first evidence that it is possible to predict the change in bone density and strength on a patient-specific basis using CT and blood markers from a single timepoint. The ability to predict these changes in bone would enable more confident decision-making when treating spinal instability. This first attempt at creating a biomarker-informed mechanobiological algorithm to predict patient-specific bone changes represents a promising step towards understanding MIBD progression and informing better clinical decision making and patient outcomes.

In conclusion, this study proves that some MM patients with vertebral lesions can be treated without surgery, proving that vertebrae in these patients do indeed rapidly strengthen in an anomalous manner. We show for the first time that this rapid remineralisation is not purely driven by healthy mechano-adaption, implying additional biological mechanisms are at play. Lastly, by training our models on blood biomarkers downstream of these signalling mechanisms, we present the possibility of generating patient-specific predictions of disease trajectory to enable further development of predictive tools to guide clinical treatments.

## Data and materials availability

All data are available in the main text or the supplementary materials.

## Supplementary Materials

Supplementary material for this article is available in the attached document.

## Dedication

The authors would like to dedicate this work to Prof. Dan Bader, who sadly passed away before its completion. Dan’s contributions to biomechanics and his commitment to applying engineering principles to clinical problems were an inspiration for this study. His work, together with his own experience of multiple myeloma, brought a deeply personal significance to our efforts to improve patient-specific prediction of fracture risk in this disease. Dan was a leading figure in bioengineering, and his legacy will endure through the people he inspired and mentored, and the pursuit of research that can make a meaningful difference to patients.

