## Supplementary Information for "Personalised prediction of anomalous bone regeneration in multiple myeloma patients"

| **Corresponding Author:** | Dr. Stefaan Verbruggen  |
| --- | --- |

**Extended Methods: Patient Data and Clinical Characteristics Description**

### Patient Cohort A: Royal National Orthopaedic Hospital (RNOH)

Clinical computed tomography (CT) data was obtained from the RNOH which included ten patients each with a baseline and follow-up scan at varying months apart (16 ±15 months), seen in Table S1.

Three different scanners were used: Phillips ingenuity (Phillips, Amsterdam, Netherlands), Phillips brilliance 64 and GE Lightspeed VCT. All protocols had the tube voltage set to 120kV while the mean tube current varied from 101-305 mA, the pixel size varied from 0.3417 mm to 0.6875 mm and slice thickness varied from 1-3 mm between protocols. The lytic vertebral level was selected by the vertebra with the largest lesion present within both baseline and follow-up CT scans.

Table S1: Patient demographics data for the RNOH MM data including age, sex, time between baseline and follow-up, vertebral level used within the scan and the condition of the vertebra.

| **Patient ID** | **Age** | **Sex** | **Time between baseline and follow-up (months)** | **Vertebral Level** | **Condition** |
| --- | --- | --- | --- | --- | --- |
| **A01** | 77 | M | 3 | T4 | Control |
|  |  |  |  | T3 | Lytic |
| **A02** | 66 | M | 12 | T10 | Control |
|  |  |  |  | T11 | Lytic |
| **A03** | 63 | M | 12 | T10 | Control |
|  |  |  |  | T11 | Lytic |
| **A04** | 74 | M | 8 | T4 | Control |
|  |  |  |  | T6 | Lytic |
| **A05** | 73 | M | 37 | T1 | Control |
|  |  |  |  | C7 | Lytic |
| **A06** | 47 | M | 2 | T4 | Control |
|  |  |  |  | T3 | Lytic |
| **A07** | 61 | M | 37 | L3 | Lytic |
| **A08** | 49 | M | 9 | T10 | Control |
|  |  |  |  | T11 | Lytic |
| **A09** | 77 | M | 38 | L3 | Control |
|  |  |  |  | L4 | Lytic |
| **A10** | 34 | M | 4 | T3 | Lytic |
|  |  |  |  | T4 | Lytic |
| **Average (±SD)** | 62 ±15 |  | 16 ±15 |  |  |

### Patient Cohort B: ANTELOPE trial

The second dataset included in this study is a time series of higher resolution quantitative CT (QCT) dataset from the ANTELOPE trial^1^, allowing quantification of bone mineral density (BMD) using an inline calibration phantom placed within the scanner. Ethical approval was obtained from the South Yorkshire Research Ethics Committee in October 2016 (IRAS ID 206171). QCT scans of the T12 vertebra were taken at baseline and 12 months after baseline. This dataset included 31 patients the androgen deprivation therapy (ADT) group, of which 29 patients completed all study assessments and 26 had a matched healthy volunteer as a control. These healthy volunteer matched controls were applied as Patient Cohort B in this study. One volunteer’s QCT scan was unusable due to overlapping vertebrae so 25 volunteers made up the cohort within this study. An overview of the healthy volunteer demographics data is shown in Table S2.

The first scans of the cohort from 2017 were taken using the GE LightSpeed VCT (GE Healthcare, Milwaukee, WI) in the radiology department at the Northern General Hospital, Sheffield, whilst the follow-up scans in 2018 along with all second scans were scanned using the Toshiba Aquilion ONE (Toshiba Medical Systems, Tokyo, Japan) at the same hospital. Quality assurance was performed once per month using a Mindways phantom (Mindways Software, Inc., Austin, TX, USA) on both scanners. The QCT protocol included a single scan from the superior edge of the T12 vertebra to the T12/L1 margin. For the GE scanner, the tube voltage was 120 kV and the mean tube current was set at 360 mA, with a voxel size of 0.937x0.937x0.625 mm^3^. For the Toshiba scanner, the tube voltage was also 120 kV, the mean tube current was set at 250 mA and a voxel size of 0.976x0.976x0.5 mm^3^.

Table S2: Patient demographics data for the ANTELOPE healthy volunteers (25 subjects) including age, height, and BMI.

| Participant ID | Age (years) | Height (cm) | BMI (kg/m^2^) |
| --- | --- | --- | --- |
| C01 | 74 | 192.3 | 31.4 |
| C02 | 76 | 169.0 | 22.3 |
| C03 | 77 | 180.1 | 30.1 |
| C04 | 73 | 173.3 | 26.2 |
| C05 | 78 | 170.4 | 26.9 |
| C06 | 68 | 182.8 | 27.1 |
| C08 | 53 | 184.4 | 26.9 |
| C09 | 63 | 188.1 | 29.6 |
| C10 | 70 | 161.0 | 29.3 |
| C11 | 79 | 175.0 | 30.7 |
| C12 | 78 | 179.0 | 32.1 |
| C14 | 74 | 168.2 | 26.8 |
| C15 | 80 | 175.2 | 26.3 |
| C16 | 82 | 169.0 | 29.7 |
| C17 | 78 | 169.0 | 24.9 |
| C19 | 75 | 180.8 | 34.9 |
| C20 | 73 | 178.0 | 24.2 |
| C21 | 78 | 159.8 | 22.7 |
| C23 | 77 | 163.2 | 31.8 |
| C24 | 73 | 184.5 | 23.9 |
| C25 | 71 | 182.0 | 24.9 |
| C26 | 71 | 173.0 | 20.4 |
| C30 | 75 | 181.0 | 25.5 |
| C31 | 71 | 171.0 | 27.6 |
| C32 | 64 | 171.7 | 23.5 |
| Average (±SD) | 73  ±6 | 175.9 ±8.0 | 27.1 ±3.5 |

### Patient Cohort C: Sheffield Teaching Hospitals (STH)

As part of the BRATS study^2^, clinical CT data at baseline and 12 months along with bone turnover marker (BTM) data (CTX, P1NP and sclerostin) at 1-, 2- and 3-months post baseline was obtained from Sheffield Teaching Hospitals with the detailed protocol published elsewhere^3^ (Table S3, Table S4). The study was conducted according to the guidelines of the Declaration of Helsinki, reviewed by the Yorkshire & The Humber – Bradford Leeds Research Ethics Committee, United Kingdom and approved by the Health Research Authority (HRA) and Health and Care Research Wales (HCRW) (REC 18/YH/0275). Two CT scanners were used; Toshiba Aquillon ONE and Toshiba Aquillon PRIME SP. Both scanners used a voltage of 120 kV and a pixel size of 1 mm. Both scanners had a slice thickness of 2 mm.

Table S3: Patient demographics data for the STH MM data including age, sex, vertebral level used within the scan, the condition of the vertebra and whether the patient had bone turnover marker (BTM) data.

| **Patient ID** | **Age** | **Sex** | **Level** | **Condition** | **BTM Data** |
| --- | --- | --- | --- | --- | --- |
| **Pt A** | 64 | M | T4 | Lytic | Yes |
| **Pt B** | 66 | M | T2 | Non-Lytic | No |
| **Pt C** | 64 | M | T8 | Non-Lytic | No |
| **Pt D** | 68 | M | T12 | Non-Lytic | Yes |
| **Pt E** | 60 | F | T12 | Lytic | Yes |
| **Pt G** | 64 | M | T12 | Non-Lytic | Yes |
| **Pt H** | 66 | F | T11 | Lytic | Yes |

| **CTX** |  |  |  |
| --- | --- | --- | --- |
| **Patient ID** | **Month 1** | **Month 2** | **Month 3** |
| **Pt A** | 0.062 | 0.033 | 0.062 |
| **Pt D** | 0.373 | 0.138 | 0.053 |
| **Pt E** | 1.386 | 0.116 | 0.07 |
| **Pt G** | 0.091 | 0.033 | 0.003 |
| **Pt H** | 0.233 | 0.033 | 0.033 |

| **P1NP** |  |  |  |
| --- | --- | --- | --- |
| **Patient ID** | **Month 1** | **Month 2** | **Month 3** |
| **Pt A** | 44.13 | 36.71 | 41.41 |
| **Pt D** | 86.86 | 54 | 4.98 |
| **Pt E** | 230 | 21.46 | 25.02 |
| **Pt G** | 17.79 | 7.84 | 8.54 |
| **Pt H** | 78.98 | 67.94 | 13.43 |

| **Sclerostin** |  |  |  |
| --- | --- | --- | --- |
| **Patient ID** | **Month 1** | **Month 2** | **Month 3** |
| **Pt A** | 15.267 | 13.568 | 19.603 |
| **Pt D** | 35.979 | 31.153 | 31.108 |
| **Pt E** | 42.285 | 33.727 | 21.444 |
| **Pt G** | 24.44 | 21.911 | 18.291 |
| **Pt H** | 29.506 | 24.44 | 26.93 |

**Extended Methods: Finite Element Modelling Set-up**

The following series of computational operations was applied to all models in all studies.

Model construction:

Each vertebra was manually segmented and reconstructed from clinical CT scans into a 3D model using 3D-Slicer (3D-Slicer 5.6.2, [https://www.slicer.org/])^4^. The vertebra models were then aligned to ensure the loading was applied perpendicular to the endplates of the vertebral body. This was conducted by creating best-fit planes on the superior and inferior endplates and aligning the vertebra to the average of these two planes using anatomical landmarks. This process was completed in Ansys SpaceClaim (Ansys®, [SpaceClaim], 2021R1, ANSYS, Inc). The endplates were also identified and labelled on the vertebral body, in SpaceClaim, to create a surface for ease of applying the boundary conditions (Figure S1). Each model was prepared using SpaceClaim to ensure sufficient quality meshing by removing anomalous sharp edges and floating elements. The models were meshed with quadratic (10 node) tetrahedral elements in Ansys Mechanical (Ansys®, [Workbench Mechanical], 2021R1, ANSYS, Inc).

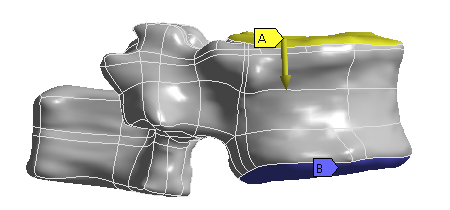

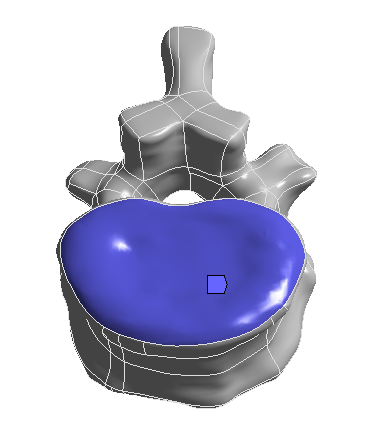

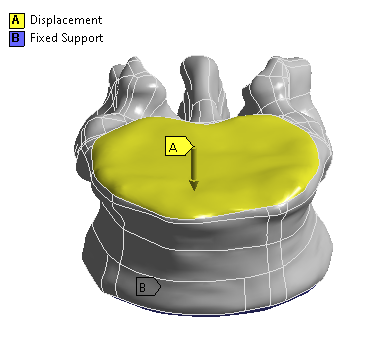

Figure S1: Application of boundary conditions. (A) The displacement applied to the superior endplate. (B) The fixed support applied to the inferior endplate.

The minimum edge size of the quadratic tetrahedral elements was set to 0.675 mm which is equal to the lowest image resolution of the CT scans. The minimum edge size was increased by a factor of 1.48 to create 3 coarser meshes (esize = 1.00, 1.48 and 2.19 mm) for mesh convergence analysis.

Material Properties:

Both bone and lytic tissue were modelled as heterogenous, isotropic, and elastic-plastic materials, following the assumption that lytic lesions only affect local bone density^5^. Bonemat (BONEMAT®, 2013) was used to map the linear material properties for each vertebra model using a patient specific densitometry phantomless calibration equation and phenomenological relationships (Equations S1 and S2), more explicitly explained in our previous study^6^. The plastic behaviour of bone was modelled as isotropic and symmetric, where the yield stress criterion is computed based on a density-strength relationship (Equation S3), and a 95% reduction in the post-yield elastic modulus (Equation S4). The density determined by CT ($\rho_{QCT}$) and ashing density ($\rho_{ash}$) is highly correlated^7^, therefore it is assumed that $\rho_{QCT}=\rho_{ash}$.

$\rho_{ash}=\rho_{QCT}=\rho_{app}\times0.6 [g/{cm}^{3}]$^7^

**Equation S1**

where $\rho_{QCT}$ is the QCT equivalent BMD and $\rho_{app}$ is the apparent density.

$E_{el}=4730{\rho_{app}}^{1.56} \left[ MPa \right]$^8^

**Equation S2**

where *E_el_* defines the elastic modulus as a function of the apparent density, $\rho_{app}$.

$\sigma_{y1}=21.7{\rho_{app}}^{1.52} [g/{cm}^{3}]$^9^

**Equation S3**

where $\sigma_{y1}$ represents the yield stress in tension as a function of the apparent density, $\rho_{app}$.

$E_{py}=0.05\times4730{\rho_{app}}^{1.56} [MPa]$^8,10^

**Equation S4**

where $E_{py}$ symbolises the post-yield elastic modulus as a function of the apparent density, $\rho_{app}$.

Boundary Conditions:

The models were loaded in compression on the superior endplate with a displacement of 1.9% of the minimum vertebral height (H) (Figure S2). The use of 1.9% apparent deformation has been experimentally proven to produce accurate ultimate loads in vertebral bodies^11–13^. The nodes on the inferior endplate were fixed in all directions (Figure S1).

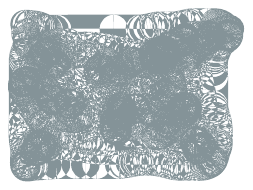

**H**

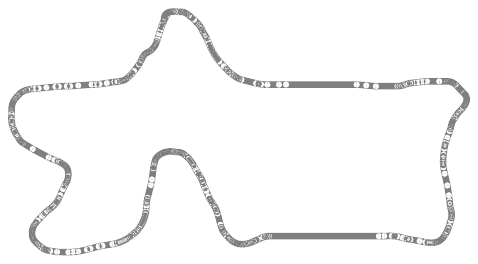

**50% H**

Figure S2: Schematic representation of the region (grey) of the vertebral bodies considered for analysis along with the definition of vertebral height (H) and 50% vertebral height.

For each vertebra level, the node containing the maximum compressive strain (EPEL3) in all models was reported. The resultant loads (F) were calculated as the sum of the axial forces on the inferior endplate of each vertebral model and displacement was computed from the axial displacement of the central node on the superior endplate. The linear range of the force-displacement curve was used to calculate the stiffness (K). The cross-sectional area (CSA) was calculated as the mean CSA of the vertebral body, excluding the endplates and posterior components. The apparent modulus (*E*) was calculated using Equation S5.

$$E=K \times\frac{H}{CSA}$$

**Equation S5**

where E is defined by spring stiffness (*K*) multiplied by the ratio between the vertebral height and CSA of each vertebra. The ultimate strength (σ_U_) is calculated using Equation S6.

$$\sigma_{U}= \frac{FL}{CSA}$$

**Equation S6**

where *σ_U_* is defined as the ratio between the resultant force at 1.9% strain (FL) and the CSA.

**Extended Methods: Verification, Validation and Uncertainty Quantification**

Intra-observer reproducibility of manual segmentation procedure

The use of manual segmentation has previously been evaluated for reproducibility^14^ and validated for FE studies of bone strength^15^. Manual segmentation is currently still the gold standard method for segmenting CT images of vertebrae with lesions, however, this process has not been investigated for reproducibility or the effect of the image quality on the segmentation, inherently affecting the prediction of mechanical properties.

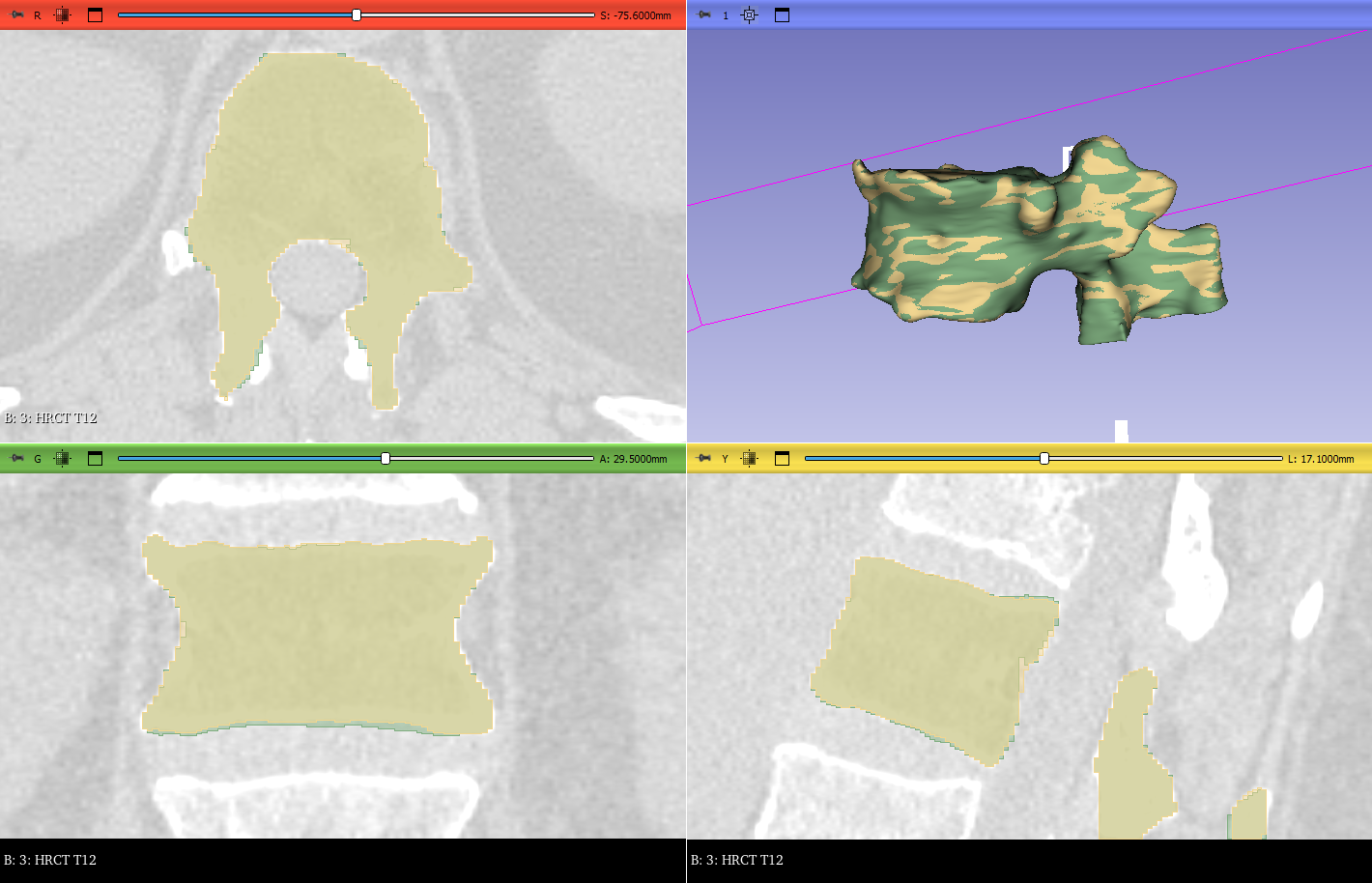

Figure S3: Example of segmentation comparison process in 3D Slicer. Two segmentations (1) in yellow and (2) in green.

In order to verify the reproducibility of this method, each vertebra with lesions from patients in Patient Cohort A was segmented three times by a skilled operator with the segmentations taking place more than 6 days apart to reduce memory effects. Masks were created for each vertebrae which represented a number of voxels defining the shape of an anatomical component^16^. Initially, thresholding was applied to define which voxels were included in the mask depending on their grey level. The threshold was adjusted by the operator until the mask was optimised for the maximum amount of bone and a well-defined cortex whilst minimising soft tissue. Once the thresholding was optimised, the mask was assessed for any bridging, areas between vertebrae where no bony tissue exists, and manually resolved to ensure only bone was included in the mask. The interior of the mask was then filled manually based on the mask boundaries in the sagittal plane. Manual detailing was necessary when the initial thresholding over or under selected the vertebrae. A final check was conducted to ensure all anatomical views of the vertebra were defined by the mask (Figure S3).

To evaluate the reproducibility of the segmentations, two evaluation metrics were selected: Dice similarity coefficient (DSC) and Hausdorff distance (HD). DSC is a good overall measure of how well something matches by assessing the overlap where a higher value indicates better agreement. HD is useful for assessing differences when there is a complex boundary which is applicable in this case as vertebrae are irregular bones^17^. HD is the maximum distance between the points on the boundary of one mask and the closest points on the boundary of the other where lower values indicate better agreement.

$$DSC= \frac{2\left| X\cap Y \right|}{\left| X \right|+\left| Y \right|}$$

**Equation S7**

where X and Y are the two masks being compared, |X ∩ Y| is the size of the overlap between X and Y and |X| and |Y| are the sizes of the masks X and Y respectively.

$$Hausdorff Distance H\left( X,Y \right) =max(h\left( X,Y \right),h\left( Y,X \right))$$

**Equation S8**

where X and Y are two sets of points on masks X and Y respectively and $h\left( X,Y \right)$ is the distance between points in X and in Y.

To assess how the image quality affects the reproducibility of the segmentation, the relationship between three parameters and the DSC and HD were evaluated: pixel size, tube current and slice thickness. Pixel size is defined by the distance between the centres of two adjacent pixels in the imaging plane. Smaller pixel size usually results in higher spatial resolution, meaning more detail can be captured in the image and lower partial volume effects. Slice thickness is the distance between the centres of adjacent slices. Smaller slice thickness provides a more continuous coverage of the scanned volume, which can improve the accuracy of volumetric measurements. Tube current is the flow of electric charge through the X-ray tube's filament, measured in milliamperes (mA). Higher tube current results in more X-rays being generated, which can improve the image quality by increasing the signal-to-noise ratio. However, increasing the tube current increases the amount of radiation delivered to the patient so this needs to be balanced to get the highest image quality without needlessly damaging the patient.

To assess the variability of using the DSC and HD the mean and standard deviation was calculated. To compare the reproducibility of the control and MM groups, a Mann-Whitney U-test was applied where significance was recognised at p < 0.05. To compare the variance in segmentation per patient between each group, Levene’s test was conducted where p < 0.05 indicated significance. The metrics were also compared to pixel size, slice thickness and tube current using a Spearman rank correlation. Due to the small sample size, an r ≤ 0.5 represents a weak correlation, 0.5 < r<0.9 is moderate correlation and 0.9 < r≤1 is a strong correlation, with p-values where significance was considered for p < 0.05. To evaluate the combined use of the image quality parameters, a multivariate linear regression analysis was performed with p-values where significance was considered for p<0.05.

Table S5 shows the overall results for the DSC, average and 95% maximum HD, pixel sizing, slice thickness and tube current for both the control and MM groups. There was no significant difference between the control and MM group for both DSC (Control: 0.97 ±0.02, MM: 0.95 ±0.03), mean HD (Control: 0.27 ±0.22 mm, MM: 0.34 ±0.22 mm) and 95% HD (Control: 0.79 ±0.38 mm, MM: 1.00 ±0.31 mm). There was a trend towards higher reproducibility for the control group, but this needs to be confirmed with a larger dataset.

**Table S4: Results overview of the DSC, average and 95% HD, pixel size, slice spacing and mean tube current for the control and MM dataset.**

|  | DSC (avg ±std) | Average HD (mm) (avg ±std) | 95% HD (mm) (avg ±std) | Pixel size (mm) (avg ±std) | Slice thickness (mm) (avg ±std) | Mean tube current (mA) (avg ±std) |
| --- | --- | --- | --- | --- | --- | --- |
| Non-lytic 1 | 0.98 ±0.009 | 0.18 ±0.09 | 0.73 ±0.37 | 0.9375 | 0.3 | 360 |
| Non-lytic 2 | 0.96 ±0.03 | 0.33 ±0.27 | 0.66 ±0.57 | 0.9375 | 0.3 | 360 |
| Non-lytic 3 | 0.95 ±0.03 | 0.43 ±0.33 | 0.97 ±0.58 | 0.9375 | 0.3 | 360 |
| Non-lytic 4 | 0.97 ±0.02 | 0.30 ±0.3 | 0.73 ±0.37 | 0.9375 | 0.3 | 360 |
| Non-lytic 5 | 0.99 ±0.004 | 0.11 ±0.03 | 0.88 ±0.11 | 0.9375 | 0.3 | 360 |
| Lytic 1 | 0.96 ±0.02 | 0.28 ±0.1 | 0.8 ±0.2 | 0.39 | 0.5 | 305 |
| Lytic 2 | 0.95 ± 0.02 | 0.41 ±0.17 | 1.38 ±0.2 | 0.67 | 1.5 | 298 |
| Lytic 3 | 0.95 ±0.02 | 0.30 ±0.13 | 0.74 ±0.19 | 0.39 | 0.5 | 186 |
| Lytic 4 | 0.92 ±0.04 | 0.60 ±0.34 | 1.22 ±0.34 | 0.69 | 1.25 | 101 |
| Lytic 5 | 0.98 ±0.003 | 0.14 ±0.013 | 0.87 ±0 | 0.34 | 0.5 | 226 |
| Non-lytic avg | 0.97 ±0.02 | 0.27 ±0.22 | 0.79 ±0.38 | 0.9375 ±0 | 0.3 ±0 | 360 ±0 |
| Lytic avg | 0.95 ±0.03 | 0.34 ±0.22 | 1.00 ±0.31 | 0.4968 ±0.15 | 0.85 ±0.45 | 216 ±70 |
| p - value | 0.095 | 0.69 | 0.2 | 0.0079 | 0.0079 | 0.0079 |
| p - value | 0.095 | 0.69 | 0.2 | 0.0079 | 0.0079 | 0.0079 |

Figure S4 shows the grouped and individual data for both evaluation metrics for the lytic and non-lytic groups. The data for the average HD was equally distributed in both the non-lytic and lytic groups (Figure S4B), as well as having a similar range. The range of DSCs and 95% HD was larger for the lytic group compared to the non-lytic control (Figure S4A & C). For the lytic group, the patients whose DSC were the highest (Lytic 1 and Lytic 5) had more similar scan properties to that of the controls (Table S5, Figure S5A) which allowed for more consistent segmentation compared to the rest of the cohort. In particular, the patient with the lowest DSC (lytic 4) had a scan of lower quality, larger pixels and slice thickness and lower tube current (Table S5, Figure S5B).

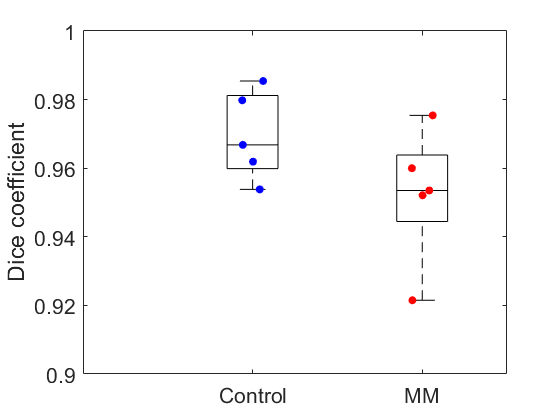

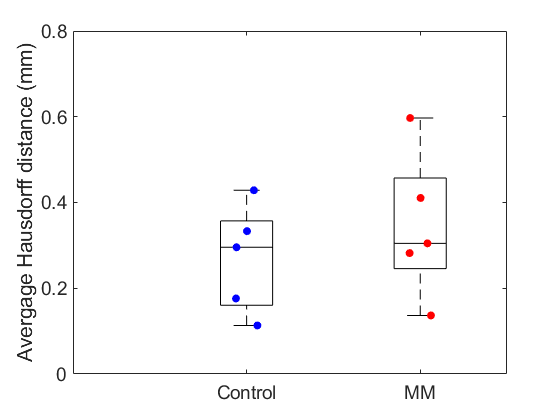

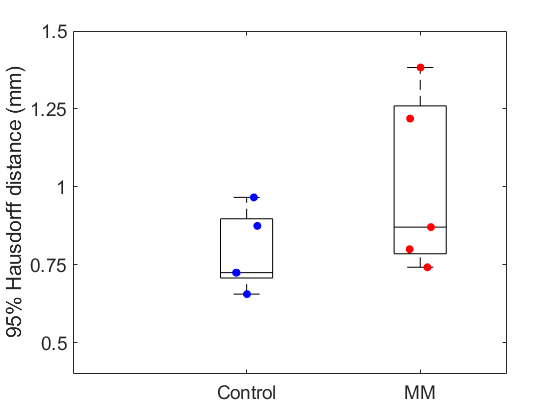

**A**

**B**

**C**

Figure S4: Box plots showing the DSC (A), average HD (B) and 95% HD (C) for the control and MM datasets.

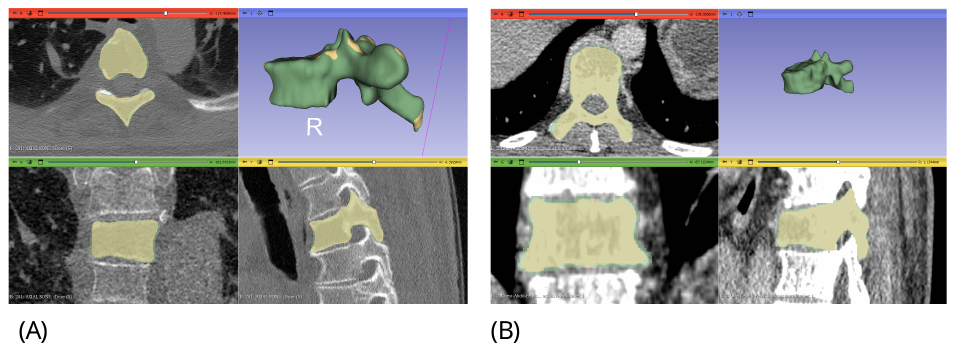

Figure S5: Representative segmentations of (A) Lytic 1 – CT scan with the best scan quality and (B) lytic 4 – CT scan with the worst scan quality.

Due to the difference in segmentation consistency in the Lytic group, the relationship between the evaluation metrics and the current, pixel size and slice thickness were evaluated. Linear regressions were constructed, and the significance of the correlation was determined (Figure S6). The current had a weak and insignificant correlation with the DSC (r = 0.5, p = 0.45) and the HD (r = -0.5, p = 0.45). However, the pixel size had a strong but insignificant correlation with the DSC (-0.9, p = 0.08) and a moderate but insignificant correlation with the HD (r = 0.8, p = 0.13). Similarly, the slice thickness had a moderate but insignificant correlation with both the DSC (r = -0.78 p = 0.2) and HD (r = 0.78, p = 0.2).

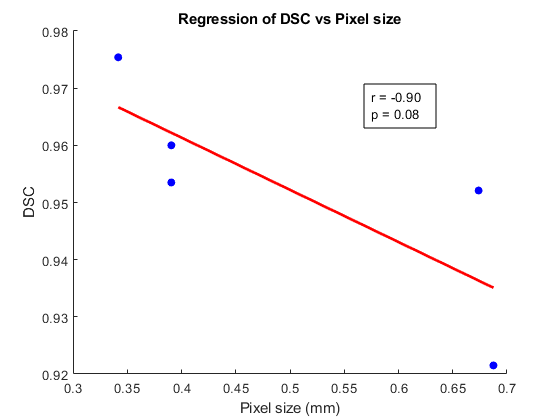

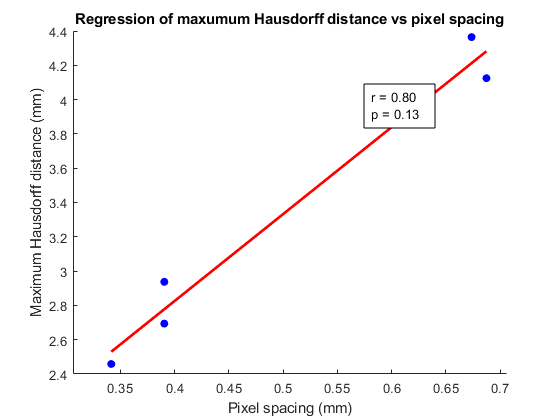

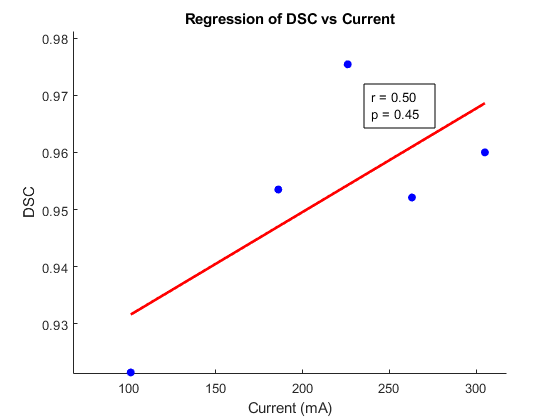

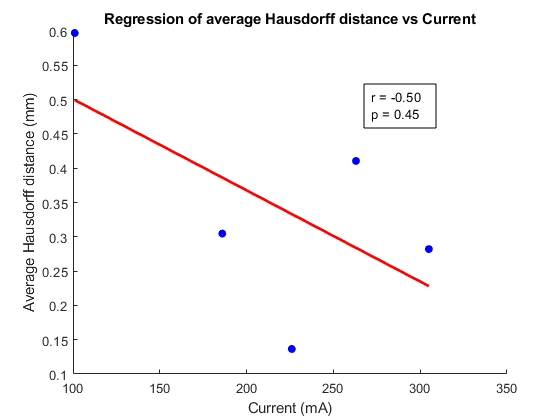

Figure S6: Linear regression analysis between (A) pixel size and DSC, (B) pixel thickness and HD, (C) tube current and DSC, (D) tube current and HD (E) slice thickness and DSC and (F) slice thickness and HD.

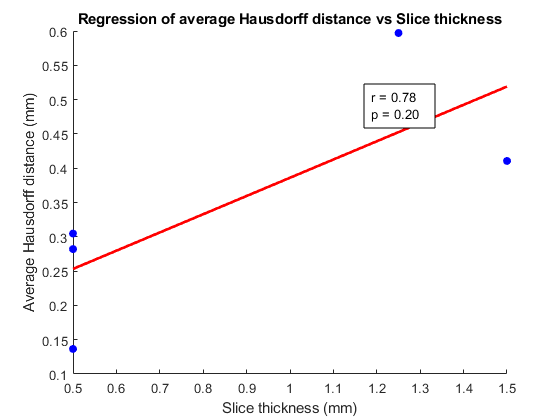

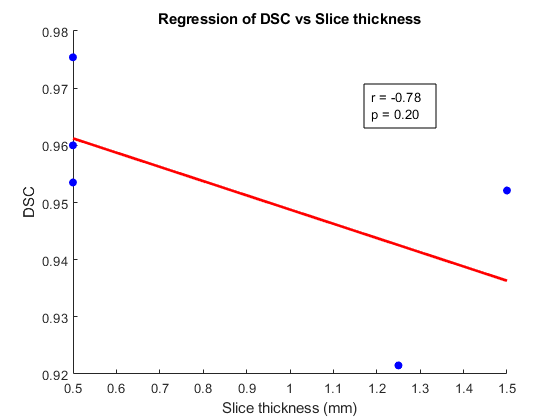

**A**

**B**

**C**
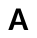

**D**

**E**

**F**

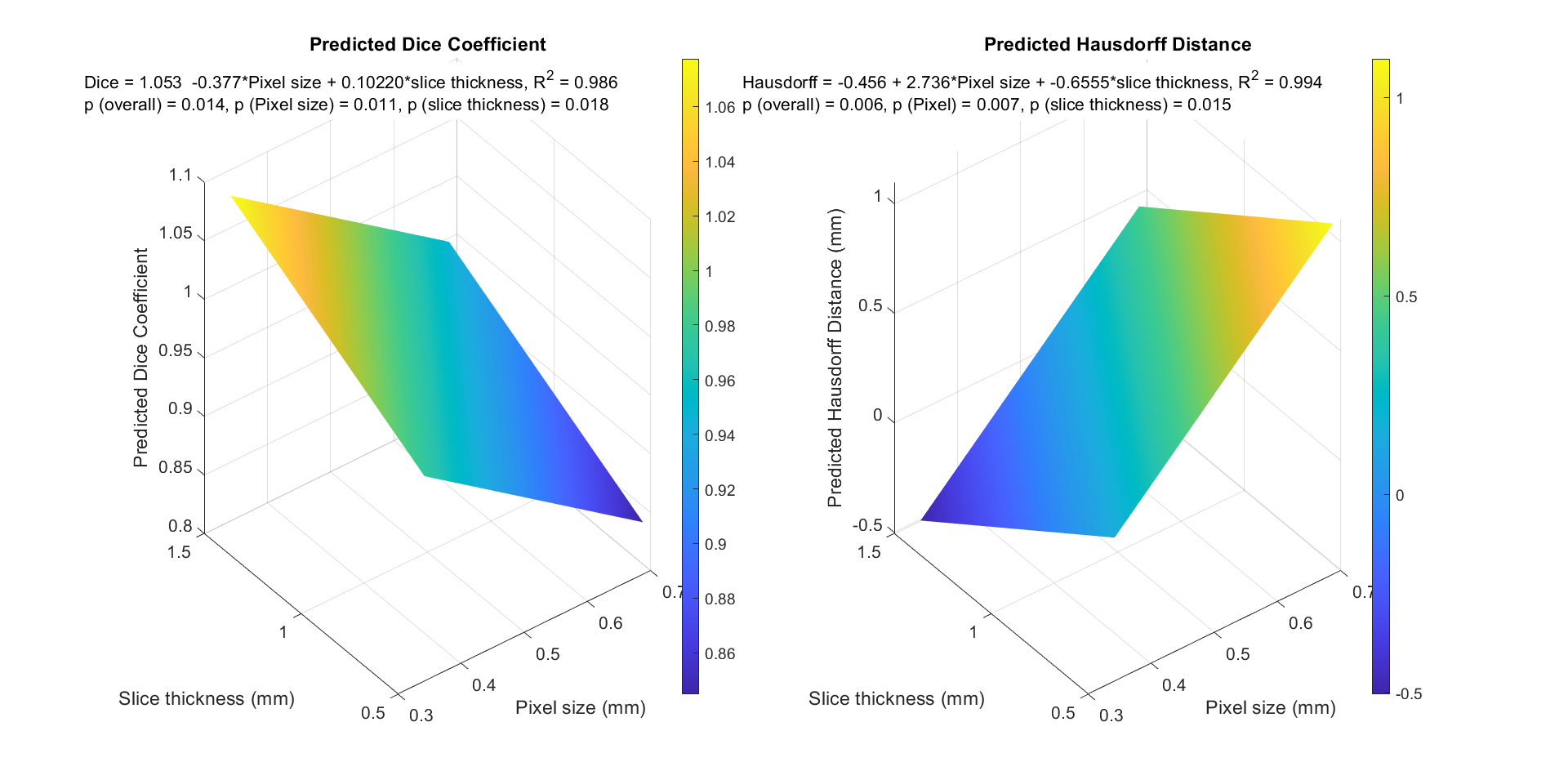

**A**

**B**

Figure S7: Multivariate analysis of the combined effects of image quality parameters and the evaluation metrics. (A) Predicted DSC from a combination of slice thickness and pixel size, (B) Predicted HD from slice thickness and pixel size.

As the relationship between the evaluation metrics and pixel size and slice thickness was moderate to strong but not significant, a multivariate analysis was conducted to understand whether the combination of these two scan properties significantly influenced the segmentation reproducibility (Figure S7). The pixel size and slice thickness could significantly (p = 0.014) explain 98.6% of the variance in the DSC. There was a significant, strong, negative relationship between pixel size and the DSC (p = 0.011) and a significant positive relationship between the slice thickness on the DSC. The pixel size and slice thickness could significantly (p = 0.006) explain 99.4% of the variance in the HD. In comparison to the DSC, pixel size had a significant, positive relationship on the HD (p = 0.007) while slice thickness had a significant, negative relationship with HD (p = 0.015).

Therefore, after this repeatability analysis we did not find significant differences between the lytic and non-lytic groups. However, we did find a significant impact of image quality on vertebral segmentation in CT scans. Pixel size and slice thickness emerged as key predictors of segmentation accuracy, with smaller pixel sizes and thinner slices leading to improved results. These findings highlight the importance of optimising image acquisition parameters to enhance the reliability and accuracy of vertebral segmentation. While these image quality parameters varied in the lytic groups’ CT scans, influencing the reproducibility, the DSC and HDs were considered acceptable for processing through the rest of the FE pipeline. For all analyses in this study, segmentations were performed once by the same trained operator to ensure robust consistency in the dataset and analyses.

Phantom and Phantomless Calibration Verification

In-line phantom calibration is currently the gold standard for calibrating CT scans and for calculating material properties of dense tissues for computational models. However, in oncology departments, it is not routine to include a calibration phantom within the scanning protocol, meaning that retrospective scans are challenging to calibrate. Therefore, in this study, we applied a phantomless calibration techniques, where the density within each scan is calibrated based on known densities captured within each scan (e.g. air). However, as this phantomless calibration has not previously been compared with the gold standard phantom calibration for vertebrae, we first conducted a verification step to ensure robust repeatability of both calibration types. To this end, we tested the phantomless method on scans from the Patient Cohort B (ANTELOPE clinical trial dataset) in which inline phantoms were included, allowing us to compare our calibration method directly with the current gold standard. This study has been published elsewhere^6^, and is briefly described for the current study below.

Briefly, all fifty QCT scans in the ANTELOPE dataset, both control and treated were analysed. The QCT scan protocol included a solid inline Mindways calibration phantom (Image Analysis, Inc., Columbia, KY, USA) containing rods of 0, 0.075, and 0.15 g/cm^3^ equivalent concentration of calcium hydroxyapatite. The densitometric calibration was computed using a standard approach, which assumes a linear relationship between the average Hounsfield units (HU) and the known equivalent mean values of equivalent BMD of each rod^18,19^.

Testing a combination of different substance densities within the scans, we found that the smallest mean BMD error value was the combination of aorta and air (0.0045 g/cm^3^). Similarly, using a Spearman’s correlation, we found that the air/aorta combination presented the strongest and most significant correlation between phantom and phantomless calibrations (r =0.98, p<0.01). This is both useful and inherently sensible, as the density of air and blood are well characterised and, relative to the density of a mineralised tissue like bone, do not vary significantly from patient to patient. Therefore, for all analyses in this study, unless otherwise stated, this phantomless calibration method (air/aorta) has been used.

Mesh Convergence Analysis

Finite element models inherently rely on the accuracy and quality of the mesh containing the individual elements. Given the intricate architecture of individual vertebrae, which have even more complex geometries when affected by cancer or dynamically adapting to mechanical load, it was important to verify the integrity of the mesh using a mesh convergence study. Taking three of the clinical-grade CT scans form Patient Cohort A (n=3 patients), three vertebrae were modelled (a lytic vertebra, and the two non-lytic lesion-free vertebrae at the spinal level above or below). For each of these vertebrae, the element sizes were increased by a factor of 1.48 (giving element sizes of 0.675 mm, 1.00 mm, 1.48 mm, and 2.19 mm), and the node containing the maximum compressive strain in all models was reported. To evaluate whether convergence was achieved, the percentage difference (%diff) was calculated with respect to the values obtained from the most refined model (element size = 0.675 mm). If the percentage difference was less than 10% for all results, the mesh was considered to be converged^20^. The distribution of the elastic modulus was also computed for each refined model and vertebra level to assess the uniformity between the most refined models.

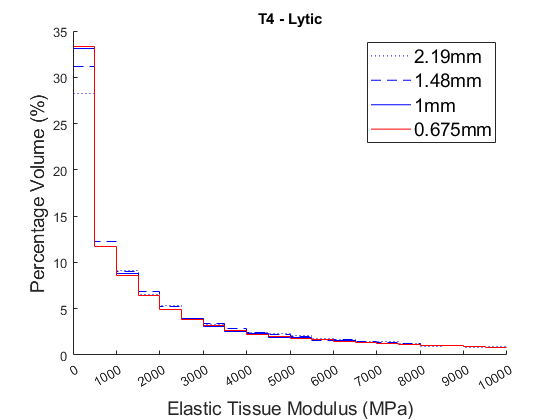

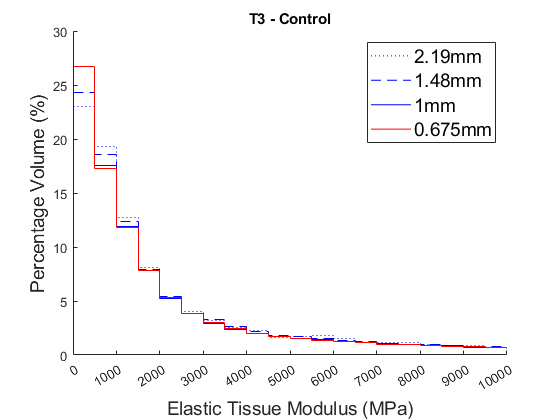

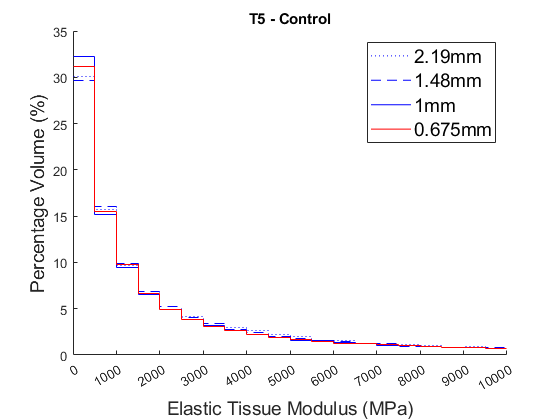

Figure S8 - Distribution of the elastic tissue modulus in each of the mesh refinement models generated from the control vertebrae (T3 and T5) and from the vertebra with the lytic lesion (T4).

The mesh convergence was assessed based on the apparent modulus (E). Changes in the elastic tissue modulus between the models with element size 0.675 mm and 1mm were minimal (Figure S8) and the decrease in element size resulted in a decrease in variability for all vertebrae. Therefore the mesh element size applied for all models in this study was 1 mm, which was also larger than the CT voxel size and the thickness of the cortical shell (<0.5 mm).

**Extended Methods: Mechanobiological algorithm**

Mechanobiological algorithm formulation

While a long history of mechanobiological algorithms exists, dating back to Frost’s Mechanostat Theory^21^, there is currently no mechanobiological algorithm describing vertebral bone remodelling in healthy volunteers or MM patients. Therefore, we developed an adaptive mechanobiological algorithm to investigate how mechanical loads affect biological processes.

The organ-level FE was coupled with an algorithm representing the cell-level behaviours and interactions of bone cells using differential equations, which adjusted material properties based on biological pathways described by differential equations. The mechanical stimulus was defined based on the difference between the SED/density (SED/ρ) from the FE model and reference stimulus from literature^22^. If the SED/ρ was above this reference value, the mechanical stimulus the element received was positive and if the SED/ρ was below this reference value, the mechanical stimulus the element received was negative. The value of the stimulus influenced the interaction between bone cells and the ratio between formation and resorption. Osteocytes weren’t explicitly modelled but their effects were indirectly considered with the osteocyte density ($d_{k}$), osteocyte network connectivity ($d_{0}$) and mechanosensitvity of the osteocytes ($u_{k}$)^22,23^.

$$S\left( t \right)=e^{({-d_{k}}/{d_{0}})}u_{k}(Sk - ({Sk}_{o} + (Sk - {Sk}_{o})(1 - e^{(-\lambda t)})))$$

**Equation S9**

where$d_{k}$= OCd/N_rem_ Osteocyte density/number of remodelling sites, $d_{0}$= osteocyte network connectivity, $S\left( t \right)$ = mechanical stimulus, $u_{k}$= osteocyte mechanosensitivity, $Sk$ =SED/ρ per element, ${Sk}_{o}$= reference SED/ρ, $\lambda$ =velocity of adaptation.

The bone cell interactions were controlled by the mechanical stimulus which influenced the number of active osteoblasts and osteoclasts. g12 represented the signal from osteoblasts to osteoclasts (e.g. TGF-β) and g21 represented the signal from osteoclasts to osteoblasts (e.g. RANKL/OPG). The method of grouping the signalling pathways into two autocrine and paracrine pathways was chosen due to the lack of data on the individual signalling pathways (e.g. RANKL/OPG)^24^.

$$\begin{matrix} g11=g22=0 \\ g12= A_{1}+B_{1}e^{-\gamma_{1}S(t)} \\ g21= A_{2}+B_{2}e^{-\gamma_{2}\left| S(t) \right|} \end{matrix}$$

**Equation S10**

where g11 and g22 represent the autocrine factors that influence osteoclasts and osteoblasts respectively. A_1_ and A_2_, B_1_ and B_2_, $\gamma_{1}$ and $\gamma_{2}$ represent the parameters influencing the paracrine factors.

The change in osteoblasts and osteoclasts was controlled by the mechanical stimulus (above) and parameters that dictate how many of the cells are active over time^25^.

$$\frac{{dx}_{C}}{dt}=\alpha_{1}x_{C}^{g11}x_{B}^{g21}-\beta_{1}x_{C}$$

$$\frac{{dx}_{B}}{dt}=\alpha_{2}x_{C}^{g12}x_{B}^{g22}-\beta_{2}x_{B}$$

**Equation S11**

where $x_{C}$ is the osteoclast population and $x_{B}$ is the osteoblast population, $\alpha_{1}$ and $\alpha_{2}$ represent the rate of formation of osteoclasts and osteoblasts respectively and $\beta_{1}$ and $\beta_{2}$ are the rates of apoptosis for osteoclasts and osteoblasts respectively.

The number of active osteoblasts and osteoclasts was then calculated by taking the steady state populations away from the updated population for that timestep (affected by the mechanical stimulus)^25^.

$$\left\{ \left. \begin{matrix} X_{C}=x_{C}-\bar{x}_{C} & ifx_{C}>\bar{x}_{C} \\ X_{C}=0 & ifx_{C}\leq\bar{x}_{C} \end{matrix} \right\} \right.$$

$$\left\{ \left. \begin{matrix} X_{B}=x_{B}-\bar{x}_{B} & ifx_{B}>\bar{x}_{B} \\ X_{B}=0 & ifx_{B}\leq\bar{x}_{B} \end{matrix} \right\} \right.$$

**Equation S12**

where $X_{C}$ and $X_{B}$ are the active populations of osteoclasts and osteoblasts, respectively. Steady state populations ($\bar{x}_{C}$ and $\bar{x}_{B}$) are calculated by setting $\frac{{dx}_{C}}{dt}=0$ and $\frac{{dx}_{B}}{dt}=0$

The change in density was then based on the number of active osteoblasts and osteoclasts and the rate of formation and resorption respectively^25^.

**Table S5: Initial Parameters for the mechanobiological algorithm**^22,25^

| **Parameter** | **Description** | **Initial Value** |
| --- | --- | --- |
| $d_{0}$ | Osteocyte network connectivity | 0.1 |
| OCd (mm^-3^) | Osteocyte density | 10625 |
| N_rem_ | Number of remodelling sites | Number of elements |
| $u_{k}$ | Osteocyte mechanosensitivity | 0.5 |
| ${Sk}_{o}$ (J/kg) | Threshold mechanical stimulus | 0.0025 |
| $\lambda$ (days^-1^) | Velocity of adaption | 0.002 |
| $A_{1}$ | Parameter to regulate production of paracrine factors in g12 | 1.6 |
| $B_{1}$ | Parameter to regulate production of paracrine factors in g12 | -0.49 |
| $\gamma_{1}$ | Parameter to regulate production of paracrine factors in g12 | 33.37 |
| $A_{2}$ | Parameter to regulate production of paracrine factors in g21 | -1.6 |
| $B_{2}$ | Parameter to regulate production of paracrine factors in g21 | 0.6 |
| $\gamma_{2}$ | Parameter to regulate production of paracrine factors in g21 | 16.67 |
| $k_{1}$ | Osteoclast resorption rate | 0.00305 |
| $\alpha_{1}$ (osteoclasts/day) | Osteoclast formation rate | 3 |
| $\beta_{1}$ (osteoclasts/day) | Osteoclast apoptosis rate | 0.2 |
| Initial population of osteoclasts | Initial population of osteoclasts | 1 |
| $k_{2}$ | Osteoblast formation rate | 0.000016 |
| $\alpha_{2}$ (osteoblasts/day) | Osteoblast formation rate | 5 |
| $\beta_{2}$ (osteoblasts/day) | Osteoblast apoptosis rate | 0.02 |
| Initial population of osteoblasts | Initial population of osteoblasts | 15 |

$$\frac{d\rho}{dt}=k_{2}X_{B}-k_{1}X_{C}$$

**Equation S13**

The density was then updated using the forward Euler method and Young’s modulus based on the relationship described earlier^8,10^.

$$\rho_{t+\Delta t}=\rho_{t}+\Delta\rho$$

**Equation S14**

$$E={4730\left( \frac{\rho}{0.6} \right)}^{1.56}$$

**Equation S15**

where $\rho_{t+\Delta t}$ is the updated density, $\rho_{t}$ is the density at the current time step and $\Delta\rho$is the change in density. *E* is the Young’s modulus and $\rho$is the density at the end of time.

In order to visualise the change in mechanical stimulus, g12, g21, active osteoclasts and active osteoblasts and then density, Figure S9 shows the evolution of these parameters over time in two randomly sampled elements in one of the vertebral models. The number of active osteoclasts was highest within the first 20-30 days and then the osteoblasts took over until around 150 days where no more remodelling occured (Figure S12). For the first random element, the initial mechanical stimulus was positive meaning there would be a net gain in bone during this remodelling cycle. This can also be seen from g12, signal from osteoclasts to osteoblasts, starting high and decreasing over time, showing the slow decrease in osteoblast differentiation due to a reduction in mechanical stimulus. Similarly, g21, signal from osteoblasts to osteoclasts slowly increasing over time, meaning an increase in osteoclast differentiation as the mechanical stimulus reduces. The impact of this on the change in density was seen by an initial decrease due to osteoclastic activity and then regaining stability through osteoblastic activity, with the final density dictated by the balance between the two.

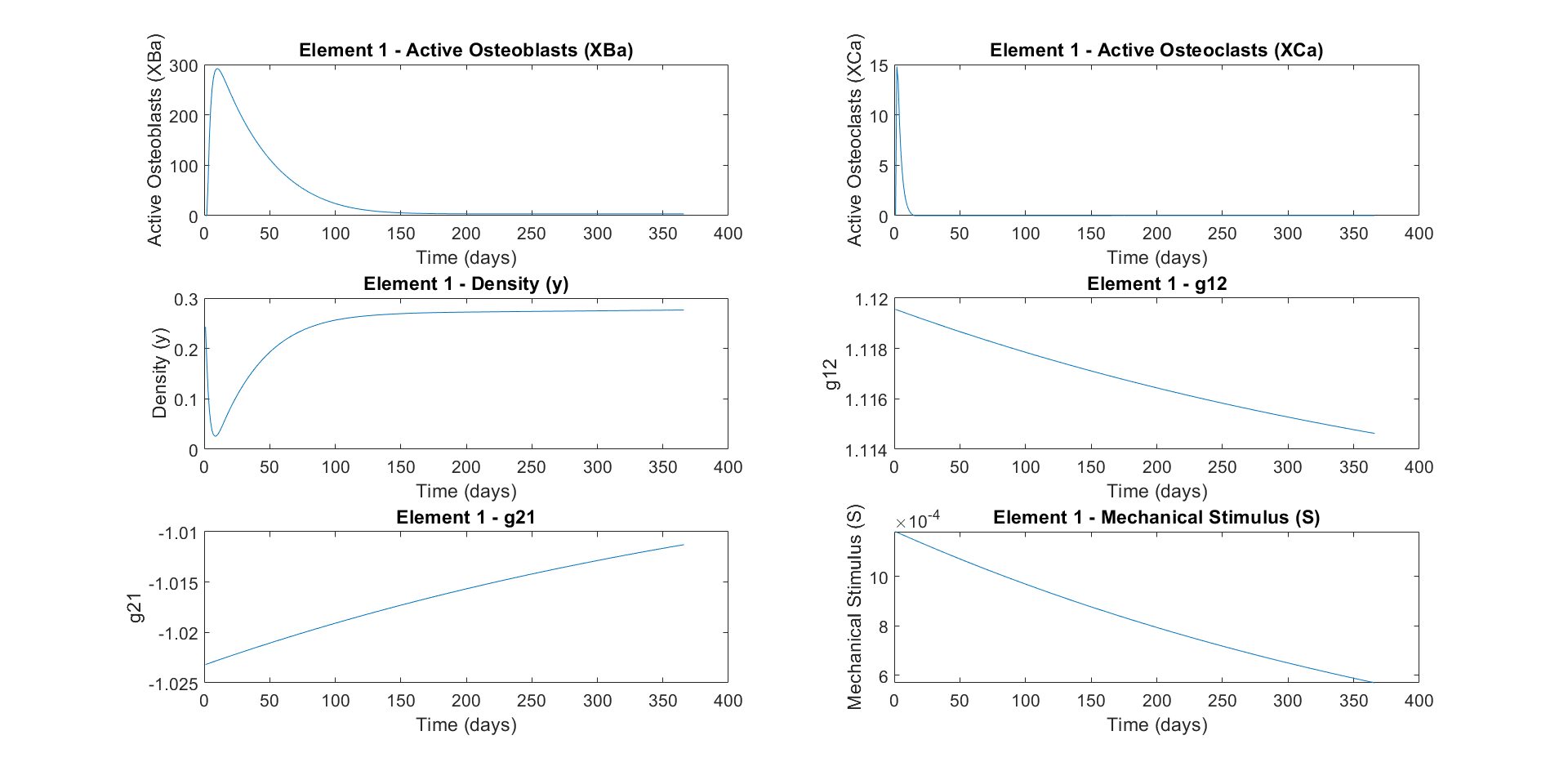

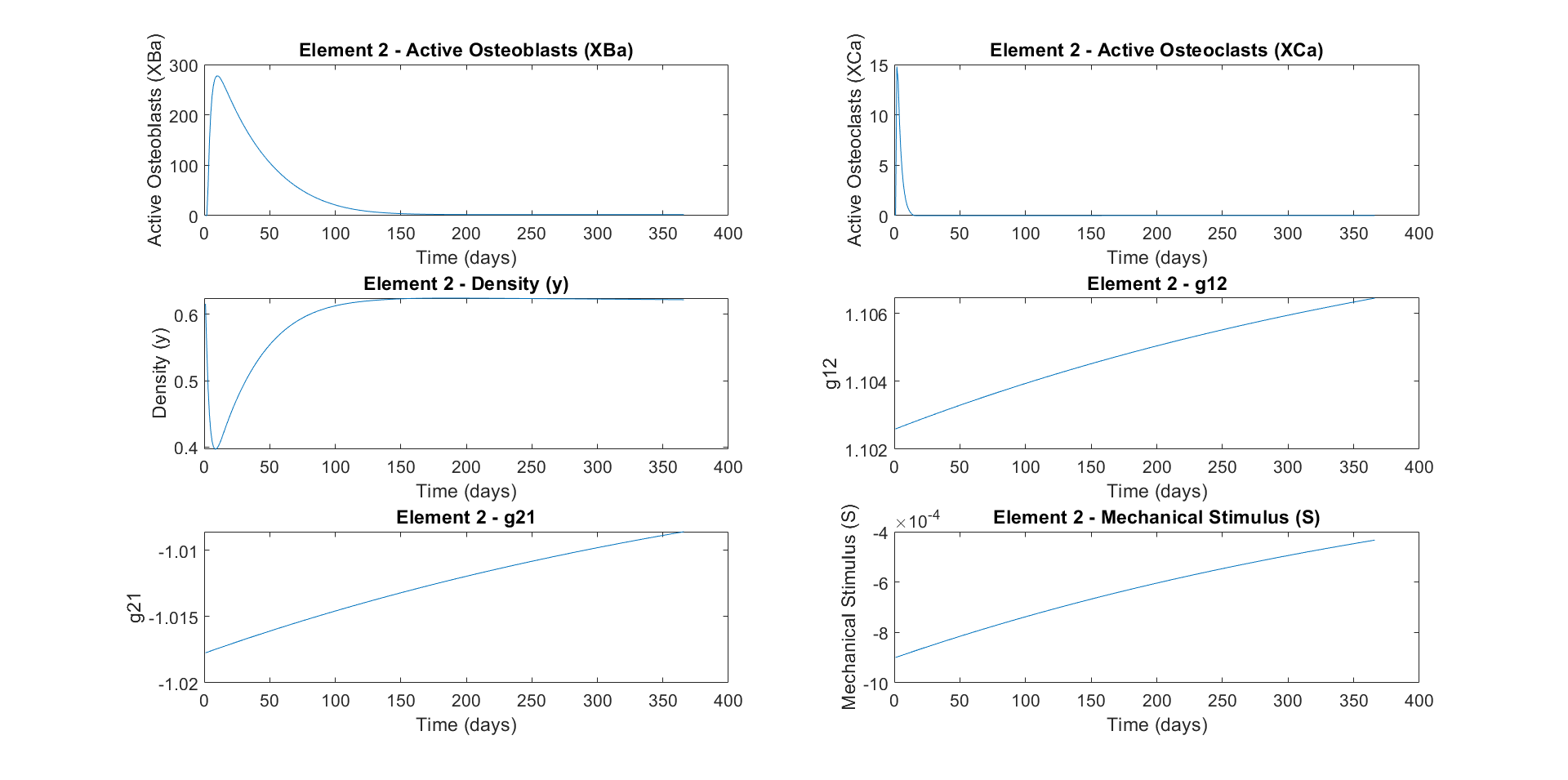

Figure S9: Example evolution of two random elements (A and B) for active osteoclasts and active osteoblasts, density, g12, g21 and mechanical stimulus.

**A**

**B**

For the second random element, the initial mechanical stimulus was negative meaning there should be a net loss in density. The positive change in g12 and g21 highlights the increase in signal from osteoclasts to osteoblasts and the signal from osteoblasts to osteoclasts. In this element, these signals ware very balanced, with the osteoblastic activity influencing the overall change more than the negative mechanical stimulus. This balance is seen from the density where it stabilises around the same point as the initial density.

Mechanobiological algorithm sensitivity and optimisation

For the healthy volunteers, a sensitivity analysis was conducted on all parameters individually to see how sensitive the algorithm was to change. Each parameter was increased and decreased by 50% and the local change in density was evaluated then averaged across all elements.

From the sensitivity, parameters were chosen to be optimised for each volunteer based on how sensitive they were. If they induced a change in density of 20-150%, they were included in the optimisation. To optimise the parameters, a MatLab script (MATLAB 24.1.0.2689473 (R2024a) Update 6) was developed which began by defining an objective function which ran the original mechanobiological algorithm, computing the simulated density and then calculating the standard error between the simulated density and the actual follow-up density.

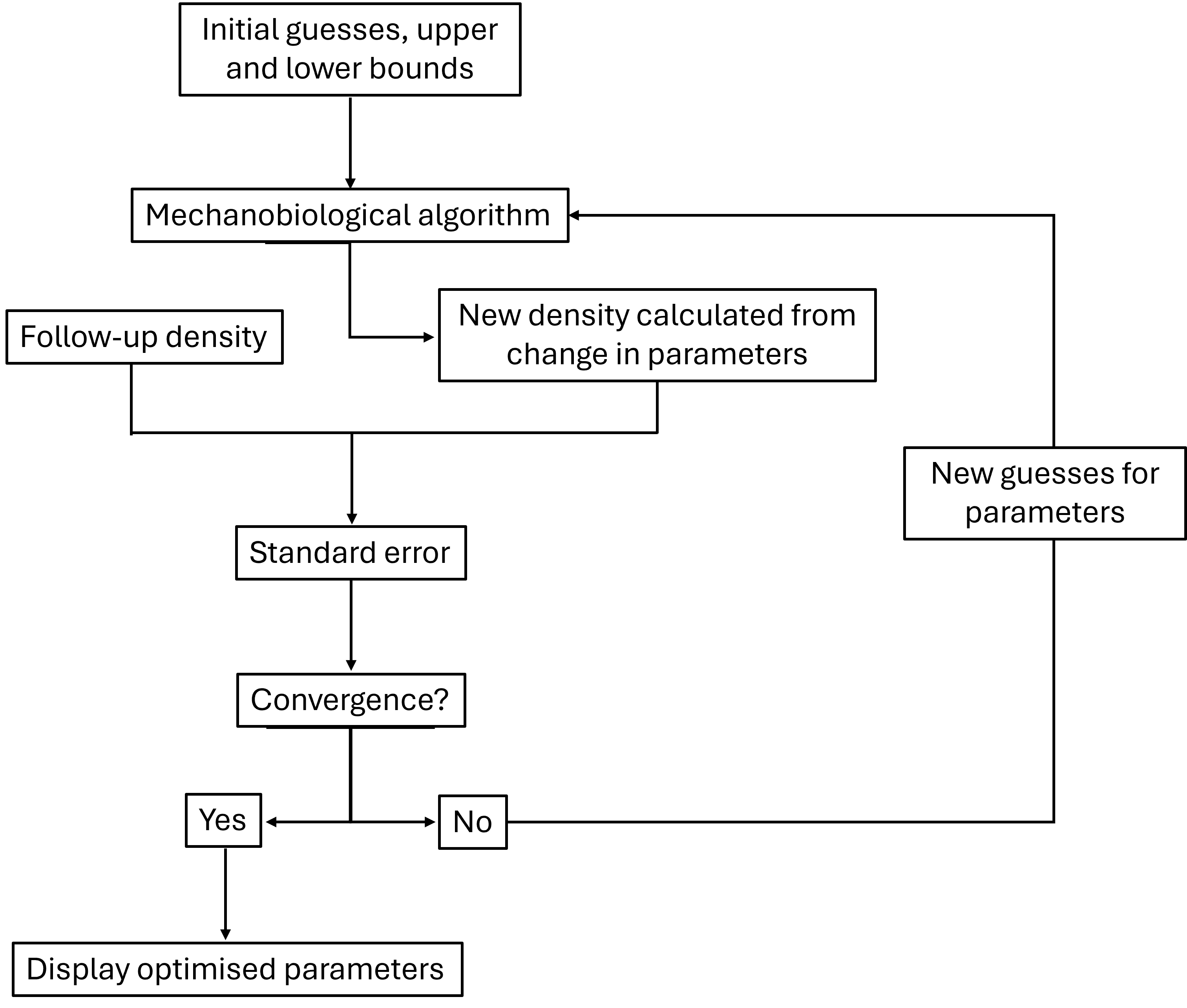

Figure S10: Flow-chart of the optimisation

The parameters to be optimised were given initial guesses (Table S5) taken from literature^22^ and upper and lower bounds which are set to ±50% of the original value. To evaluate the results of each set of parameters, the function fmincon was used. An options structure for the fmincon function was created to specify how the optimisation algorithm should behave including the display level (iterative) and the algorithm to use sequential quadratic programming (SQP) which means each iteration of the optimisation was solved and stored and with each iteration, the SQP refines its approximation to get closer to the optimal solution). The fmincon function was then called together with the objective function, initial guesses, bounds, and options. This function performs the optimization to minimise the standard error by adjusting the parameters until convergence. Convergence was met when either the optimality tolerance, the standard error was below $1 \times{10}^{-6}$, or the step tolerance was below $1 \times{10}^{-10}$, where further changes in the input parameters did not significantly reduce standard error.

To compare the optimised parameters between the healthy volunteers, MM patient non-lytic and MM lytic groups, Mann-Whitney Unpaired test was used, where significance was defined as p < 0.05. To test for significance (p < 0.05) between the follow-up BMD and FL and the predicted BMD and FL, a Wilcoxon paired test was used. Linear regression analyses were performed between the follow-up and predicted BMD and FL to assess the variance. A significant result was found when p < 0.05.

Figure S11 shows the average percentage change in density in all elements when increasing and decreasing each parameter by 50%. It is clear that some parameters are less sensitive, with a change in density less than 10% despite a 50% increase or decrease of the input parameters (d_0_, d_k_, u_k_, λ, Sk_o_, γ_1_ and γ_2_). Osteocyte network connectivity (d_0_) and osteocyte density (d_k_) are used in Equation S9. This exponential was always going to be between 0 and 1, meaning any changes in d and d_k_ do not alter the overall response. Osteocyte mechanosensitivity (u_k_) started off very low as it was introduced as a dimmer switch between 0 and 1 and was set to 0.5. This value was too small for this parameter to increase or decrease the mechanical stimulus and density by more than 10%. The adaption velocity (λ) dictated how fast the system responds to the mechanical changes. This parameter should have a low sensitivity as bone remodelling occurs over a long period meaning the effect of λ is spread across the time. If bone remodelling was a shorter process (days instead of months), the effect of λ would be more apparent. The difference in mechanical signal from changes in Sk_o_ were not large enough to affect the osteoblast and osteoclast populations which drove the change in density, hence why Sk_o_ had a low sensitivity (<10%). γ_1_ and γ_2_ were exponential decay parameters that determined the influence of the mechanical stimulus S on the rates of osteoblast and osteoclast differentiation (g12 and g21). These parameters sat within Equation S9 as $e^{-\gamma_{1}S(t)}$ and $e^{-\gamma_{2}\left| S(t) \right|}$, so doubling gamma still gave a very similar value to the initial estimate.

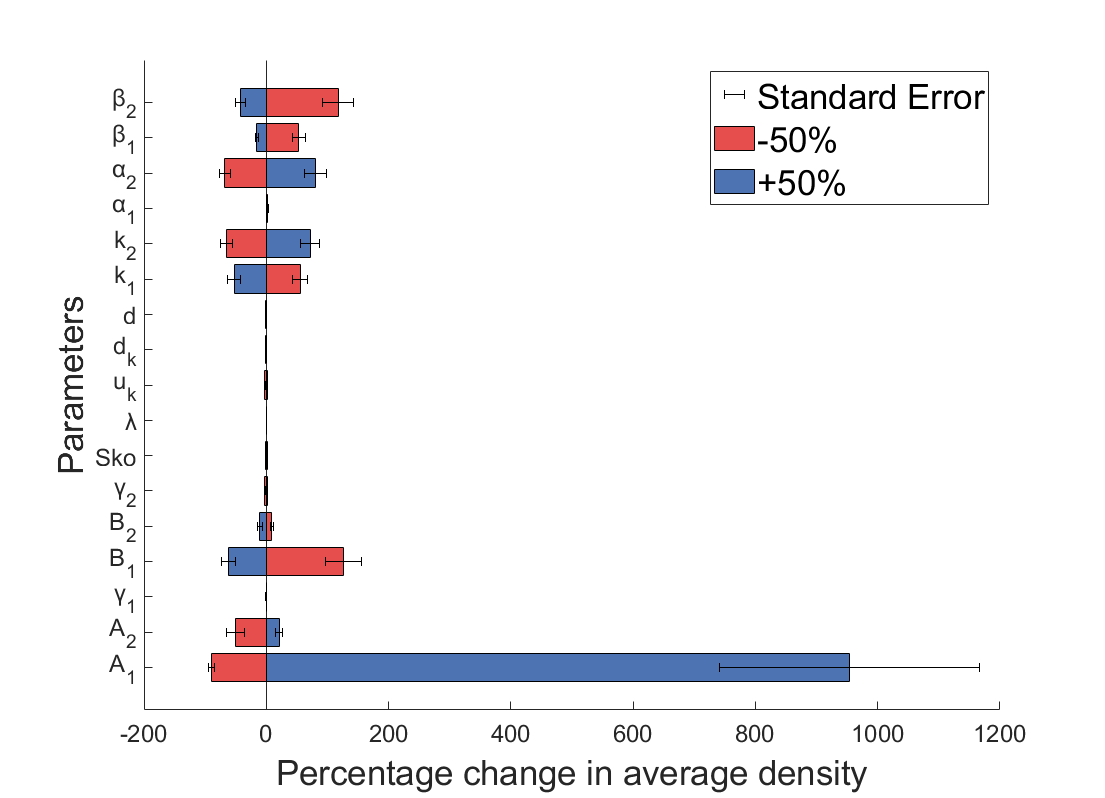

Figure S11: Change in average density when increasing and decreasing each parameter by 50% to understand the sensitivity of each parameter

It is also noticeable that A_1_ was very sensitive to a positive change (+50% in A_1_). A_1_ was the dominant factor in g12 as the exponential element was always close to 1. Equation S11 then used g12 as an exponential to the osteoclast population ( $x_{C}^{g12}$). As A_1_ was the dominant factor in g12, if A_1_ was doubled, $x_{C}^{g12}$ would increase by a factor of 76, also increasing the population of active osteoblasts by a factor of 76 for every iteration. This induced a ‘runaway’ effect where the density of each element was then significantly increased, hence the large variation in the positive change in A_1_. This did not happen for the decrease in A_1_ as the system was osteoblast dominant and therefore could recover from a decrease in osteoblast recruitment. A_2_ also did not have this effect, which was most likely due to the initial osteoblast number being 1 and therefore there was not the ‘runaway’ effect we observed with A_1_.

Figure S12 shows a zoomed in version of the sensitivity results for k_1_, k_2_, $\alpha_{1}$, $\alpha_{2}$, $\beta_{1}$ and $\beta_{2}$. Most of these parameters had ~50% change in density, both when increasing and decreasing the parameter by 50%, as would be expected for a balanced algorithm. The only parameter which seems to have little effect is $\alpha_{1}$. $\alpha_{1}$ influenced the number of osteoclasts produced per timestep but the dominant factor in this equation was most likely $x_{B}^{g21}$ as the number of initial osteoblasts at each timestep was much larger than the number of osteoclasts. Therefore, the effect of $\alpha_{1}$ was dampened. All six of these parameters have important links to biological processes that can be monitored in clinic via blood BTMs and could be altered depending on patient specific data. $\alpha_{1}$ and $\alpha_{2}$ control the osteoclast and osteoblast differentiation rate, $\beta_{1}$ and $\beta_{2}$ control the osteoclast and osteoblast apoptosis rate and k_1_ and k_2_ control the rate of resorption and formation. For this reason, these six parameters were chosen for optimisation.

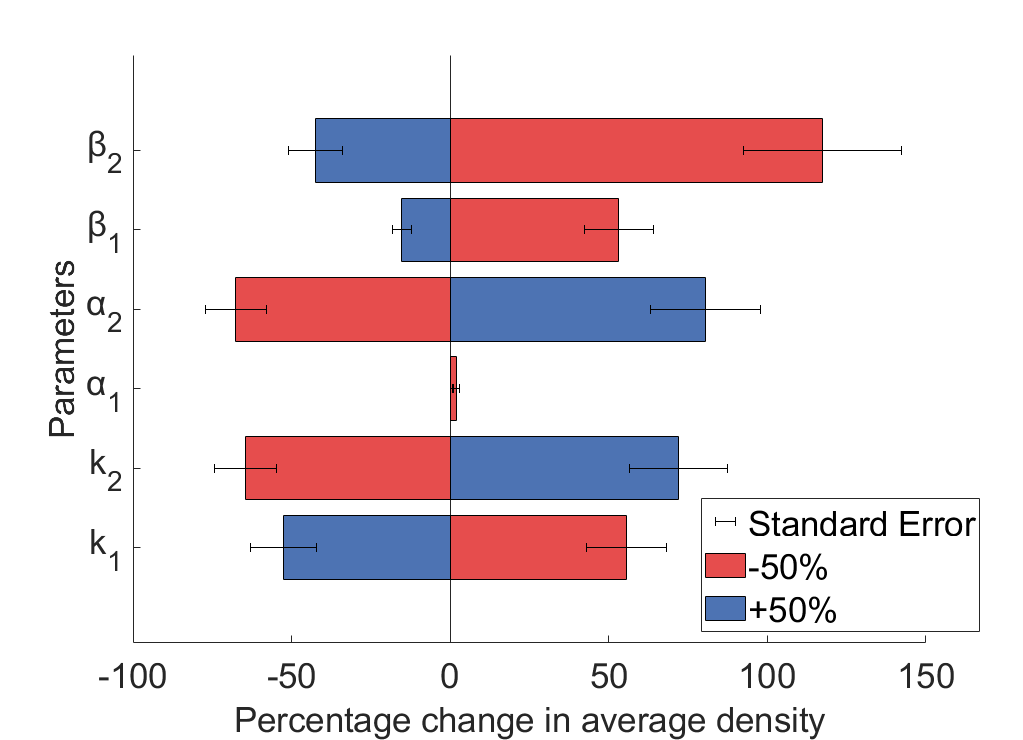

Figure S12: Close up view of the average change in density of k1, k2, $\boldsymbol{\alpha}_{\boldsymbol{1}}$, $\boldsymbol{\alpha}_{\boldsymbol{2}}$, $\boldsymbol{\beta}_{\boldsymbol{1}}$ and $\boldsymbol{\beta}_{\boldsymbol{2}}$when increasing and decreasing each parameter by 50%.

When optimising the mechanobiological parameters, both k1 and k2 decreased when moving from healthy volunteers to MM patients non-lytic vertebraes and again to the MM lytic vertebrae (Figure S13). Despite this difference being insignificant, this suggests that in MM vertebra with lesions, both bone formation and resorption rates were altered, with k_1_ decreasing more sharply than k_2_. The sharp decline in k_1_ suggested reduced bone resorption in the MM lesion group, contributing to the increased bone apposition post treatment. The decrease in k_1_ and k_2_ suggested the rate of turnover is slower in the MM patients but whether formation and resorption were balanced or not was unclear from this figure. Therefore, the ratio between k_1_ and k_2_ was calculated to understand if there was an imbalance in formation and resorption. Figure S14 shows the ratio of k_1_ to k_2_ in each group, with it decreasing slightly between healthy volunteers to the MM non-lytic group and again to the MM lytic group. This difference was not significant, but it showed there was a trend towards more osteoblastic activity compared to osteoclastic activity in MM patients and particularly the vertebrae with lesions.

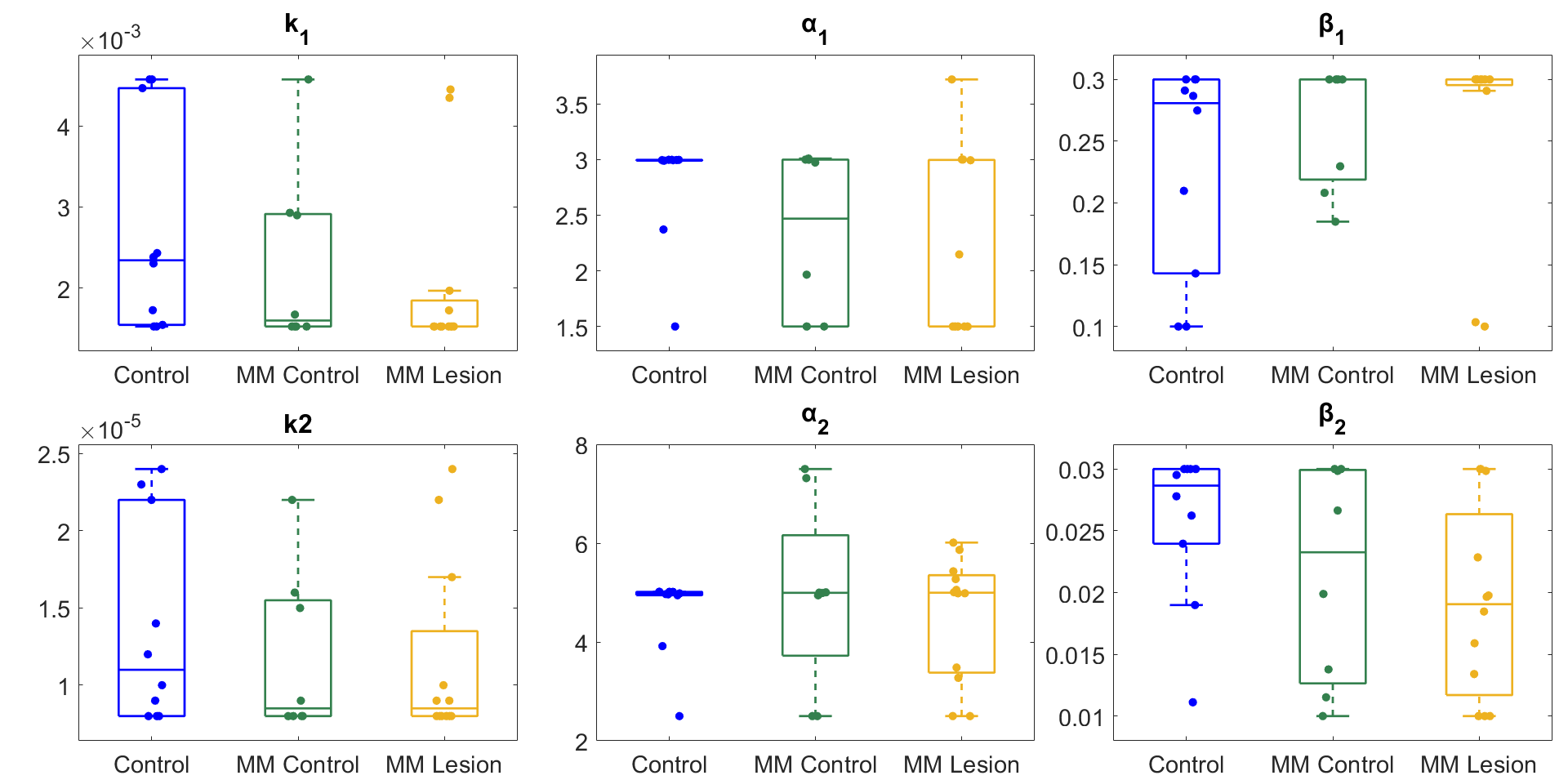

Figure S13: Box plots of optimised parameters in the control, MM control and MM lesion groups for (A) k1 and k2 (B) $\boldsymbol{\alpha}_{\boldsymbol{1}}$ and $\boldsymbol{\alpha}_{\boldsymbol{2}}$ and (C) $\boldsymbol{\beta}_{\boldsymbol{1}}$ and $\boldsymbol{\beta}_{\boldsymbol{2}}$.

**A**

**B**

**C**

For the differentiation rates, $\alpha_{1}$ showed a decrease from healthy volunteers to MM groups while $\alpha_{2}$ showed a very similar median value for all three groups. For both $\alpha_{1}$ and $\alpha_{2}$, the healthy volunteer group had a much smaller range than the other two groups, suggesting these parameters could be fixed for the controls. When looking at the ratio between $\alpha_{1}$ and $\alpha_{2}$, the MM lytic group had a much lower ratio than the controls. This suggests that the number of osteoblasts differentiating was higher than osteoclasts in the MM lytic group, however this was not significant.

For the apoptosis rates, $\beta_{1}$ was relatively stable across the groups, with no significant differences, but showed a slight increase in MM non-lytic and MM lytic, indicating that osteoclasts were being removed quicker. $\beta_{2}$ showed more variability but tended to decrease in the MM lytic group, suggesting that osteoblasts were being removed at a slower rate. However, with reduced osteoblast production this could balance overall osteoblast activity. The ratio between $\beta_{1}$ and $\beta_{2}$ had a small increase from healthy volunteer to MM non-lytic and a large jump to the MM lytic subgroup. This suggested the rate of osteoclast apoptosis was faster than osteoblasts apoptosis meaning there were more active osteoblasts.

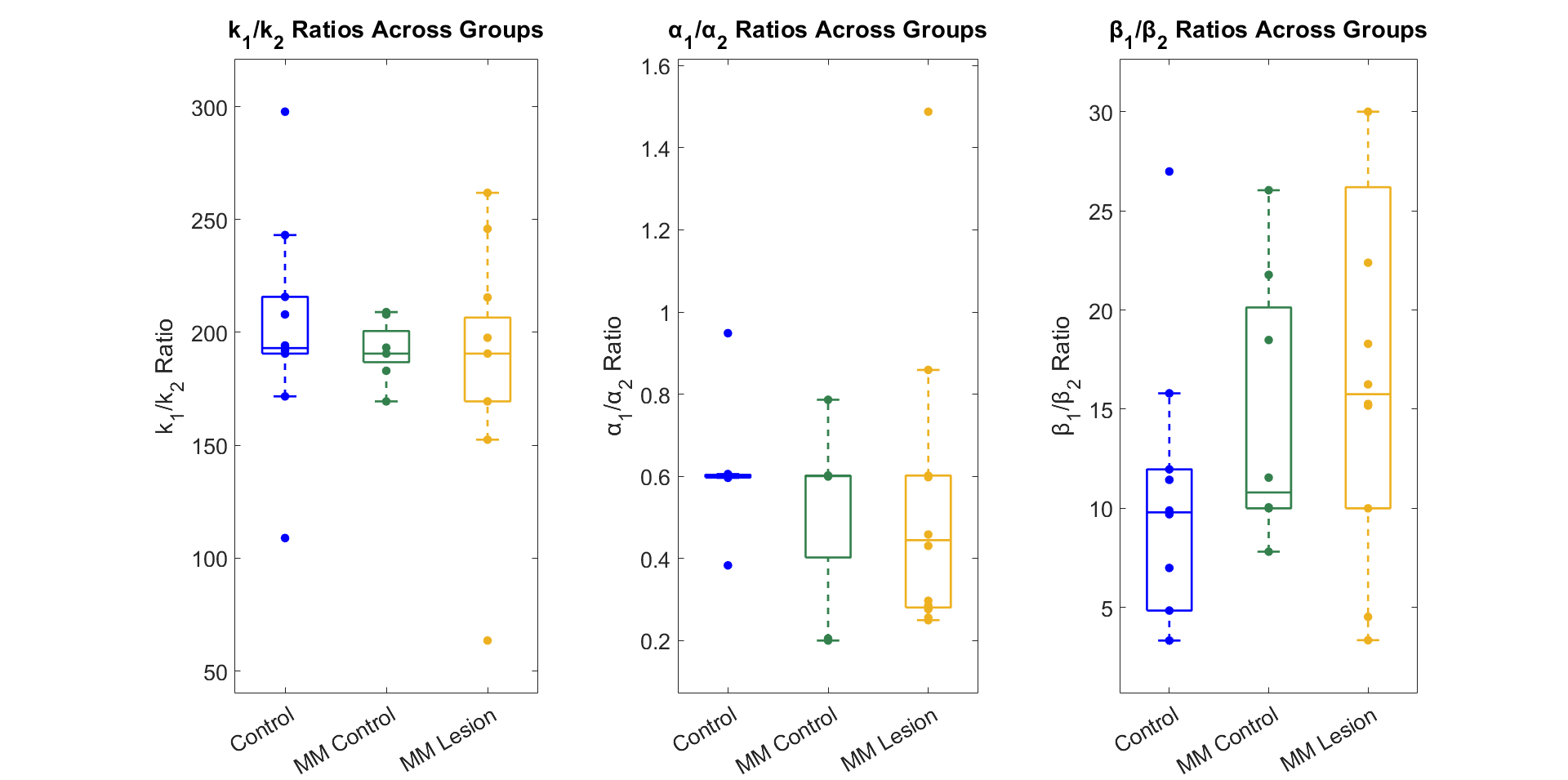

Figure S14: Ratio of (A) resorption to formation (k_1_/k_2_) (B) formation of osteoclasts to formation of osteoblasts ($\boldsymbol{\alpha}_{\boldsymbol{1}}$/$\boldsymbol{\alpha}_{\boldsymbol{2}}$) and (C) apoptosis rate of osteoclasts to apoptosis rate of osteoblasts ($\boldsymbol{\beta}_{\boldsymbol{1}}$/ $\boldsymbol{\beta}_{\boldsymbol{2}}$).

**A**

**B**

**C**

When assessing the differentiation to apoptosis ratio (Figure S15), osteoclasts had a lower ratio in the MM lytic group compared to the control and MM non-lytic, suggesting there was more osteoclast apoptosis than differentiation, suggesting less osteoclastic activity. On the other hand, the $\alpha_{2}$/$\beta_{2}$ ratio increased through the groups, suggesting more differentiation compared to apoptosis of osteoblasts, and hence more bone apposition in the MM lytic group.

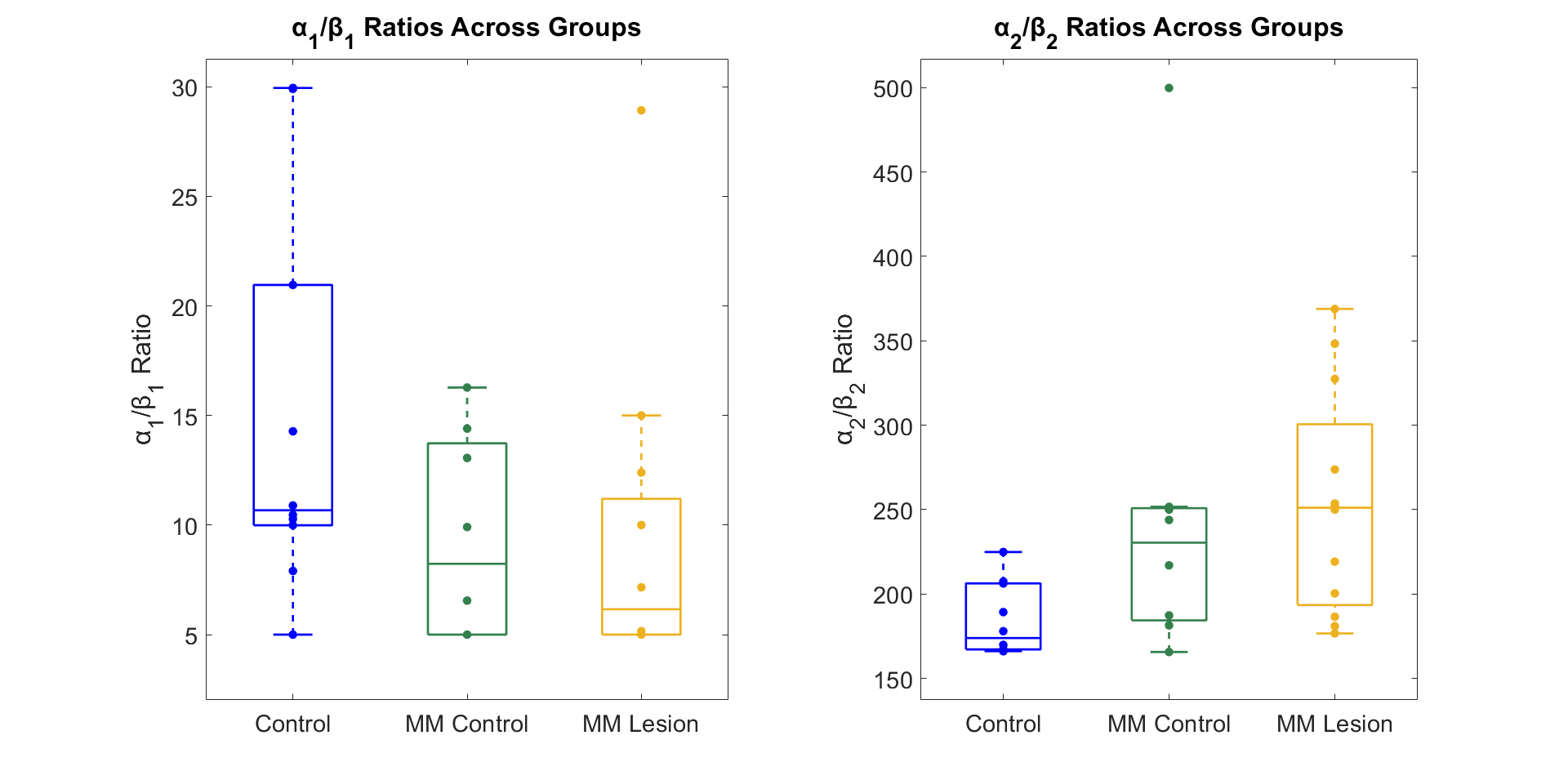

Figure S15: Ratio of (A) osteoclast formation to osteoclast apoptosis ($\boldsymbol{\alpha}_{\boldsymbol{1}}$/$\boldsymbol{\beta}_{\boldsymbol{1}}$) (B) ratio of osteoblast formation to osteoblast apoptosis ($\boldsymbol{\alpha}_{\boldsymbol{2}}$/$\boldsymbol{\beta}_{\boldsymbol{2}}$)

**A**

**B**

**Extended Methods: Clinical bone biomarker integration in mechanobiological algorithm**

The previously optimised mechanobiological algorithm was applied to the dataset from Patient Cohort C, optimising the six parameters (k1, k2, α_1_, α_2_, β_1_, β_2_) for each patient. Then, a multiple linear regression analysis using leave one out cross validation (LOOCV)^26^ was performed in MATLAB (R2024a). The multiple linear regression assessed the relationships between the three biomarkers and each of the optimised parameters (k1, k2, α_1_, α_2_, β_1_, β_2_), as well as between the three biomarkers and both BMD and failure load at 1, 2, and 3 months after baseline, separately. LOOCV was chosen due to the very small sample size, and despite its tendency to over predict outcomes ^27^, it is the best choice for this study as no other methods are viable at this scale. LOOCV used all but one patient to train the algorithm and then tested on the final patient, increasing the training dataset for small cohort sizes^26^. It repeated this for all patients, increasing the robustness of the relationship compared to a simple linear regression. Following this analysis, parameters with a strong positive correlation between follow-up and predicted (r > 0.8) were then estimated using the linear regression equations utilising the biomarkers (Equation S16). All other parameters that had a negative, moderate or weak correlation (-1 < r < 0.8) were averaged across the cohort.

$${P= \varphi_{0}^{P}}+\varphi_{CTX}^{P}\cdot CTX+\varphi_{P1NP}^{P}\cdot P1NP+\varphi_{Sclerostin}^{P}\cdot Sclerostin$$

**Equation S16**

where, *P* is the parameter to be predicted (k1, k2, α_1_, α_2_, β_1_, β_2_) , $\varphi_{0}^{P}$ is the intercept, $\varphi_{CTX}^{P}$ is the proportionality coefficient for CTX, $\varphi_{P1NP}^{P}$ is the proportionality coefficient for P1NP, $\varphi_{Sclerostin}^{P}$ is the proportionality coefficient for Sclerostin.
